# Comparative snRNAseq study of C9orf72, SOD1, and sALS spinal cord

**DOI:** 10.1101/2025.08.26.672029

**Authors:** Shaolong Cao, Mark Sheehan, Victor Cox, Dirk M. Walther, Amanda J. Guise, Edward D. Plowey, Maria I. Zavodszky, Thomas Carlile, Ravi Challa, Yi Chen, Guoqiang Zhang, Helen Mclaughlin, Catherine L. Worth, Chia-Yen Chen, Shih-Ching Lo, William Wei-Lun Chen, Jessica A. Hurt, Dann Huh

## Abstract

Amyotrophic lateral sclerosis (ALS) is a devastating neurodegenerative disease characterized by the loss of motor neurons, yet the cell-type specific molecular alterations within the spinal cords are not well characterized. In this study, we conducted deep molecular profiling of spinal cord tissues donated by people living with sporadic, C9orf72, and SOD1-ALS using single-nucleus RNA sequencing (snRNAseq). We observed numerous distinct gene expression patterns and enriched pathways among ALS types. However, when focusing on common features, we identified activation of stress-response and inflammatory pathways in specific microglia subtypes, as well as disrupted vesicle transport and synaptic function in a ventral inhibitory neuronal subtype. Notably, CPLX3, a SNARE regulator, was uniquely expressed in alpha-motor neurons and was commonly downregulated across all ALS types. While this study uncovers the molecular heterogeneity underlying ALS, it also highlights shared pathways within specific cell types, especially in the ventral inhibitory neurons that have been less explored in ALS research.

## Main Text

Approximately 5-10% of people living with ALS have familial history of the ALS, of these, ∼ 76% have a known genetic component^1^. The remaining 90-95% are classified as sporadic ALS (sALS)^2^. A repeat expansion in the C9orf72 locus (“C9”) is the most common known genetic cause of ALS (∼34% of familial cases^2^), and exhibit TDP43 pathology as in sALS. In contrast, pathogenic mutations in the SOD1 gene (“SOD1”) account for approximately 2% of ALS cases and do not display this pathology^3^. To deepen our understanding of molecular characteristics of ALS, and to identify features common across multiple types of ALS per cell type toward potential therapeutics, we conducted snRNAseq on spinal cord tissues from people living with sALS, C9, and SOD1 and non-neurodegenerative controls, as well as RNAseq and proteomics on spinal cords from different anatomical regions of sALS and controls.

### RNAseq and proteomics of sALS

We performed RNAseq on four regions – dorsal horn (DH), ventral horn (VH), ventral white matter (VWM), and whole spinal cord (WSC) – of lumbar spinal cords from sALS and controls (“cohort1”) (Fig. 1a). The VWM exhibited the highest number of differentially expressed genes (DEG), followed by VH (Fig. 1b and c), in agreement with a previous spatial transcriptomics study^4^. At pathway level, neuroinflammatory signatures were notably activated across all regions (Fig. 1e), consistent with a large-scale transcriptomics study^5^. Given that large numbers of DEGs (nDEG) were observed in VWM and VH, we performed proteomics study on these two regions. Unlike the transcriptomics, nDEG were similar between these regions (Fig. 1b and c). Notably, there were relatively few common DEGs between the RNAseq and the proteomics data (Fig. 1c), as observed in other studies on neurodegenerative diseases^6,7^. At pathway level, proteomics showed stronger dysregulation of Kinesins Dysregulation and Mitochondrial Dysfunction but weaker neuroinflammation signatures compared to transcriptomics (Fig. 1e). Also, RNA splicing pathways were activated in both VH and VWH (Supplementary Table 3). This observation led us to examine differentially spliced genes (DSG) from the RNAseq data (Method). The VH had the greatest number of DSGs (Fig. 1b), potentially reflecting TDP-43 mislocalization-induced aberrant splicing or degeneration of cells within this region. The enriched pathways of DSGs showed limited overlap across the anatomical regions (Fig. S1), nor were there large overlap between DEGs and DSGs (Fig. S2). Notably, UNC13A was identified as a DSG in VH (Fig. 1d), not due to the disease-associated cryptic exon between exons 20 and 21^8^, but due to a change in the inclusion of exons 36 and 38 (Supplementary Table 4).

**Fig. 1.**
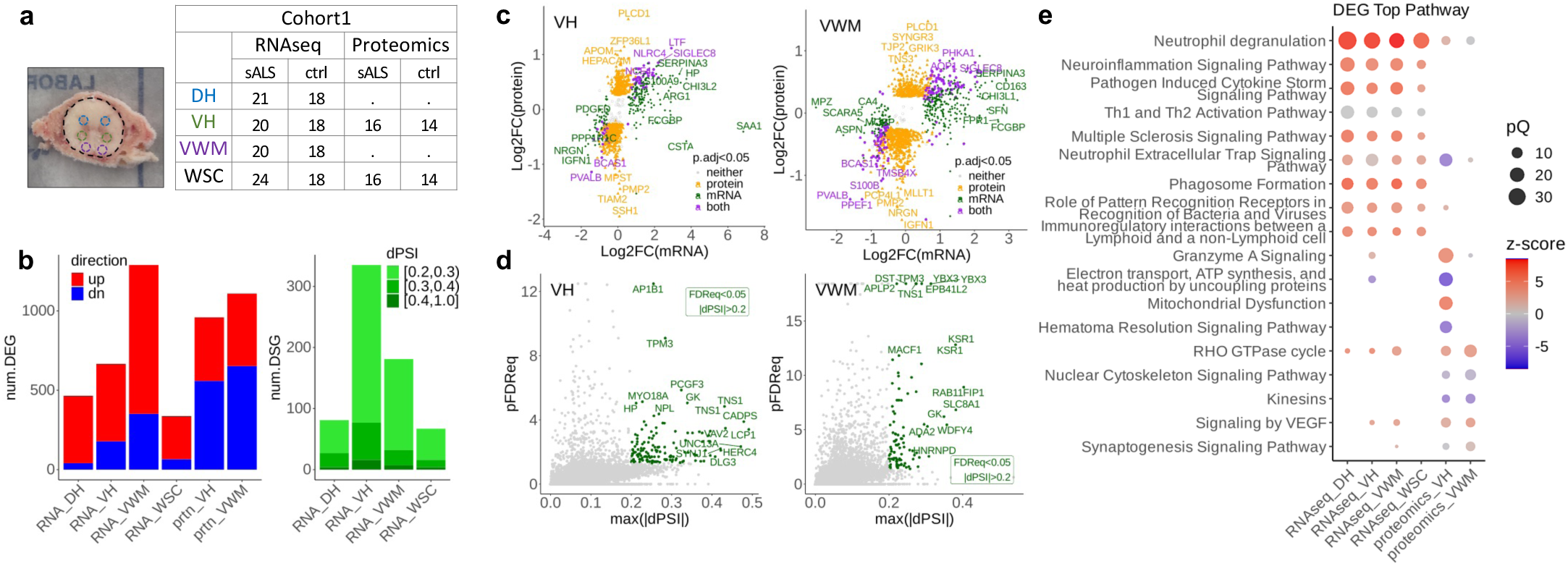
Transcriptomic and proteomic changes in sALS tissues. (a) Number of tissues for each region subjected to RNAseq or proteomics. (b) Number of DEG (p.adj<0.05 and |Log2FC|>1.2 throughout the paper) and DSG (genes with any splicing event with FDR<0.05 and |dPSI|>0.2 for rMATs and Leafcutter, and P(dPSI)>0.95 and |E(dPSI)|>0.2 for Majiq). “up” and “dn” denoting up- and down-regulated genes, respectively. (c) Log2FC of DEG from RNAseq and proteomics for VH and VWM. (d) Volcano plot of DSG for VH and VWM, where FDReq=FDR for rMATs and Leafcutter, and 1-P(dPSI) for Majiq. pFDReq is –Log10(pFDReq), and throughout the paper “pX” denotes – Log10(X), analogous to pH or pKa. Max(|dPSI|) is the maximum absolute dPSI per each gene. (e) Top 5 enriched pathways per tissue and modality from DEGs are plot together, using Ingenuity Pathway Analysis (IPA), redundancy mitigated (Method). Q denotes adjusted p-value, and positive and negative z-scores correspond to activation and inactivation of the pathway.

TDP43 aggregation, observed in 97% of people living with ALS^9^, causes both toxic gain and loss of functions. Hence, we inspected correlation between TDP43 pathology and mRNA or protein levels. Only a few genes met the DEG threshold (Fig. S3), likely due to limited sample size (14). To our knowledge, none of these genes have been previously associated with TDP-43 pathology, nor did they overlap with the DSG.

Overall, transcriptomics and proteomics show severe molecular changes occurring in the ventral spinal cords in sALS.

### snRNAseq of sporadic, C9, and SOD1 ALS spinal cords

To identify transcriptional changes specific to different cell types in ALS, we conducted snRNAseq on lumbar spinal cords from two cohorts – cohort1: sALS and control as described in the RNAseq study, and cohort2: C9, SOD1, and control. To examine ventral regions and to enrich for motor neurons, we isolated nuclei separately from VH and from the remainder of the spinal cords (WSC-VH), aiming to recover equal amounts of nuclei from each region (Fig. 2a, Method), yielding total of 773,198 nuclei from 85 subjects.

**Fig. 2.**
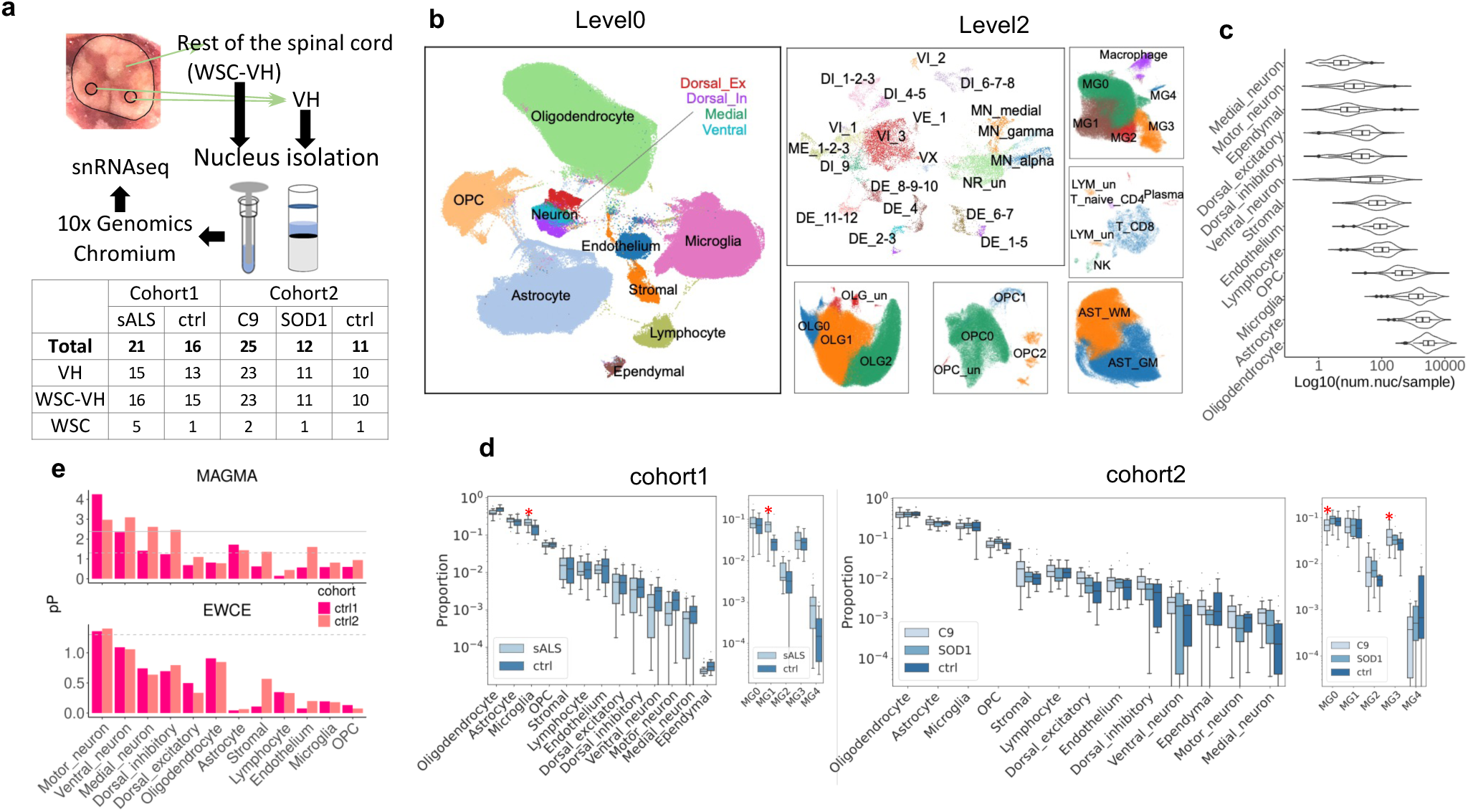
Single-nucleus RNAseq of ALS spinal cords. (a) Cartoon of 1:1 sampling of nuclei form VH and the rest of the region (WSC-VH), and number of samples per region per ALS type and control. When VH punch was not feasible, WSC was process. Note that there are two cohorts, processed separately. (b) Umap of snRNAseq with cell-type annotations, at high-level (level0) and subtype-level (level2). (c) Violin plot of number of nuclei detected in each sample per cell type. (d) Cell type proportional change from scCODA for cohort1 and cohort2. All cells in level0 and microglia level2 types are shown. Asterisk (*) denotes Prob>0.9 and 25% or more changes in cell fraction compared to respective controls. (No cell type passed this criteria in cohort2 level0). (e) Enrichment of expression of genes from ALS GWAS, using MAGMA cell typing and EWCE. Expressions in control samples from cohort1 and 2 are used (ctrl1, ctrl2), P is nominal p-value, dotted and solid lines represent P=0.05 and Q=0.05, respectively.

In this data, we classified the nuclei into 12 broad cell types (“level0”, Fig. 2b), then further clustered them into 41 subtypes (“level2”). Note that “ventral” neurons are not confined to ventral regions^10^ (Fig. S6). Despite the VH enrichment strategy, motor neurons were still rare with median of 13 nuclei per subject (Fig. 2c), which was consistent with an expectation from a post-hoc analysis (Method). Nonetheless, we identified three motor neuron subtypes: Alpha-, gamma-, and “MN_medial”, which expressed high levels of SCG2, a marker of motor neurons from medial motor column^11^ (Fig. S7e). For level2 microglia, we applied GO enrichment on the de-novo clusters for their characterization^12^, and also compared with previously reported subtypes^10,13–16^ (Fig. S9). Signatures of disease-associated microglia were not specific to a single cluster^14^ (Fig. S7d). In contrast, microglia subtypes found in Alzheimer’s disease^12,13^ were present in ALS - MG0: homeostatic, MG1: autophagy/cell migration/endocytosis, MG2: ribosomal/iron metabolism, MG3: heat-shock/hypoxia, and MG4: proliferating. This commonality implies conserved functions of microglia between ALS and AD even in different regions.

We next investigated differences in cell-type composition between ALS and controls. We downsampled the number of nuclei from VH prior to remixing with those from WSC-VH (Method), to compensate for its over-representation within the sample. Using scCODA^17^, we found that the proportion of microglia was increased in sALS, due to the increase of MG1 (Fig. 2d). While this agrees with a previous report^5^, nuanced difference was observed in cohort2, where slight increase of MG3 was observed at the expense of MG0 (Fig. 2d).

Genetically associated genes to ALS have been reported to have higher expression in neuronal cells from brain cortex^18^. Using the snRNAseq data from this study, both MAGMA cell typing^19^ and EWCE^20^ further refined that the gene sets have high expression in motor neurons, followed by ventral neurons (Fig. 2e). These results corroborate that ALS is a motor neuron disease, and also suggests potential role of ventral neurons in ALS.

### Cell-type specific transcriptomic differences among ALS types

We next investigated cell-type specific DEGs for each ALS type compared to controls. Ventral neurons consistently exhibited high nDEG driven by VI_3 population while VI_1/2 had few nDEGs (Fig. 3a), and motor neurons showed moderate nDEG. Microglia had a relatively modest nDEG, especially considering its large population size (Fig. 2c) and noticeable neuroinflammatory signatures in the RNAseq study (Fig. 1e). In following sections, we focus on ventral neurons for their prominence in DEGs, motor neurons for their relation to ALS etiology, and microglia for their implication in neuroimmunology. (Refer to Fig. S7-10).

**Fig. 3.**
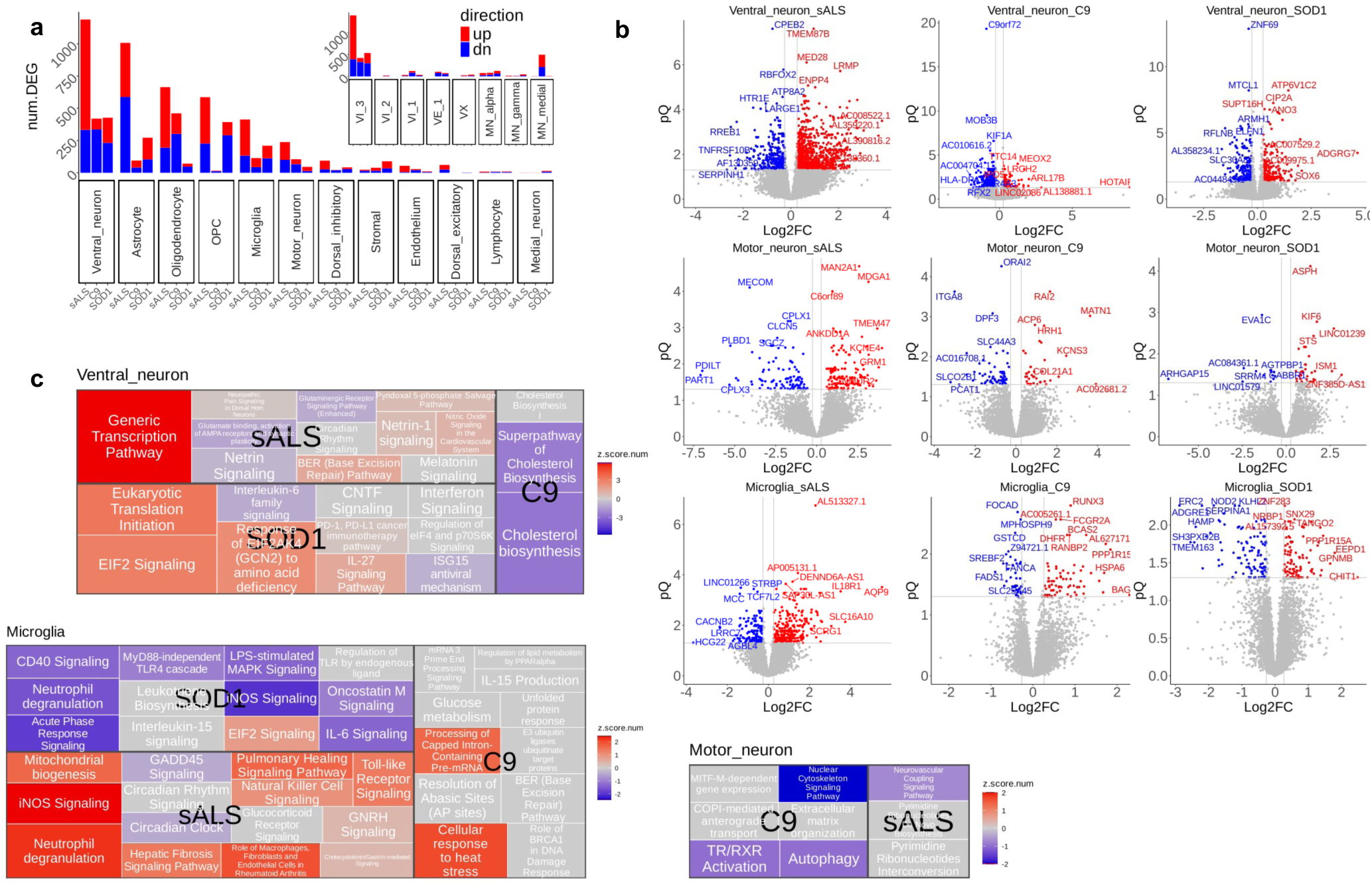
DEG and pathway analysis of ventral neuron, motor neuron, and microglia from snRNAseq. (a) number of DEG per level0 cell type, and that of ventral neuron and motor neuron level2 types (inset). (b) Volcano plots of ventral neuron, motor neuron, and micrdoglia per ALS type. Q is adjusted p-value. (c) Top enriched pathways from DEGs using Ingenuity Pathway Analysis (IPA), redundancy mitigated (Method). The size of the boxes are proportional to pQ, and positive and negative z-scores correspond to activation and inactivation of the pathway. Treemap plot is used instead of dot plots as overlaps across ALS types were few.

The top DEGs and enriched pathways seldom overlapped across ALS types (Fig. 3b and c, Fig. S11). In sALS ventral neurons, dysregulation was observed in generic transcription, netrin signaling, and neurotransmitter binding. In C9, there was a strong inactivation of cholesterol biosynthesis pathway, suggesting the abnormal cholesterol levels observed in CSF of C9-ALS variants carriers^21,22^ might reflect metabolic changes in ventral neurons. In SOD1 ALS, translation and neuroinflammation were among the highest dysregulated pathways.

We next investigated pathways enriched in motor neurons, whose degeneration is the most prominent characteristic of ALS. C9 motor neurons showed several enriched pathways, including dysregulation of Extracellular Matrix Organization as observed in Krach *et al*^23^. There also were disrupted autophagy with down-regulation of ATG9B and SQSTM1^24^ (Supplementary Table 6). In sALS, Pyrimidine Ribonucleotides Metabolism was enriched, which impairs membranes and synapse formation^25^. SOD1 motor neurons did not show any significant pathway enrichment. MN_alpha – the most vulnerable cells in ALS – showed dysregulation in synaptic functions (sALS, SOD1) or cell-cell junctions (C9, SOD1) (Fig. S14). The cryptic exon of STMN2 was upregulated in C9 MN_alpha, but it did not reach statistical significance in sALS (Supplementary Table 5).

In microglia, pathway enrichment also varied across ALS types. This goes against our hypothesis that microglial response would be similar regardless of the mechanism of motor neuron death. In sALS, Neutrophil Degranulation and TLR signaling were activated (Fig. 3c), consistent with the RNAseq study. In SOD1, however, many of pathways from sALS were also enrich but in opposing directions. In addition, inflammation-related proteins elevated in sALS CSF or serum, such as CHIT1^32^ and GPNMB^33^, were specifically expressed in microglia and also up-regulated in the SOD1 microglia but not in the sALS microglia (Fig. 3b). In C9, top pathways were related to stress response driven by heat-shock proteins (Fig. 3c).

As in the RNAseq study, we correlated the sALS snRNAseq data to TDP43 pathology scores (Fig. S15). Stromal cells showed the largest nDEG, followed by ventral neurons. Notably, in sALS ventral neurons, cholesterol biosynthesis pathways were inactivated (Fig. S15c), as in C9 (Fig. 3c). For microglia at level0, impairment of iron homeostasis was correlated with TDP43 pathology (Fig. S12c), which is due to transcriptional dysregulation rather than MG2 proliferation, as MG2 fraction did not increase with TDP43 pathology (Supplementary Table 7).

In the RNAseq study of sALS, the enrichment of neuroinflammatory signatures (Fig. 1e) implies substantial involvement microglia. To test this, we performed EWCE and found that gene sets from enriched pathways and from up-regulated genes in RNAseq study, were highly expressed in microglia (Fig. S16a). We also re-ran DEG on the RNAseq with and without microglial fraction estimated from snRNAseq as a covariate. The DEGs were similar (Fig. S16b), indicating the effect of transcriptional changes outweigh proliferation. In contrast, EWCE with DSG showed that DSG were highly expressed in motor and ventral neurons (Fig. S16d), suggesting motor and ventral neurons were the source of the high DSGs in VH (Fig. 1b), hence reinforcing the relevance of these neurons.

Together, DEG and pathway analysis highlight diversity of transcriptional response across ALS types. Despite such differences, we next explored conserved features among the ALS types, as they all share common clinical manifestations, and would be expected to share underlying molecular signatures.

### Common transcriptomic features across ALS types

Common DEGs across ALS types were mostly found in VI_3 population (Fig. 4a). Among the top genes, TMEM106B is known to function in lysosomal transport, and is also genetically linked to dementia^26,27^. The risk variant increases TMEM106B expression^28^, which was also observed in VI_3 (Fig. 4a). BICDL1, a dynein adaptor protein, was downregulated, potentially impairing retrograde transport. This is reminiscent of the splice variant of KIF5A that hyperactivates anterograde transport^29^. In addition, these observations are consistent with the downregulation of kinesins in the sALS proteomics study (Fig. 1e). In MN_alpha, CPLX3, a complexin that regulates synaptic vesicle fusion akin to UNC13A, was downregulated across all ALS types (Fig. 4a), specifically expressed in MN_alpha (Fig. S17). It was also part of SNARE and Synaptogenesis pathways enriched in sALS and SOD1, respectively (Supplementary Table 6). Taken together, these common DEGs show the importance of impaired axonal transport and synaptic functions in ALS in ventral and motor neurons.

**Fig. 4.**
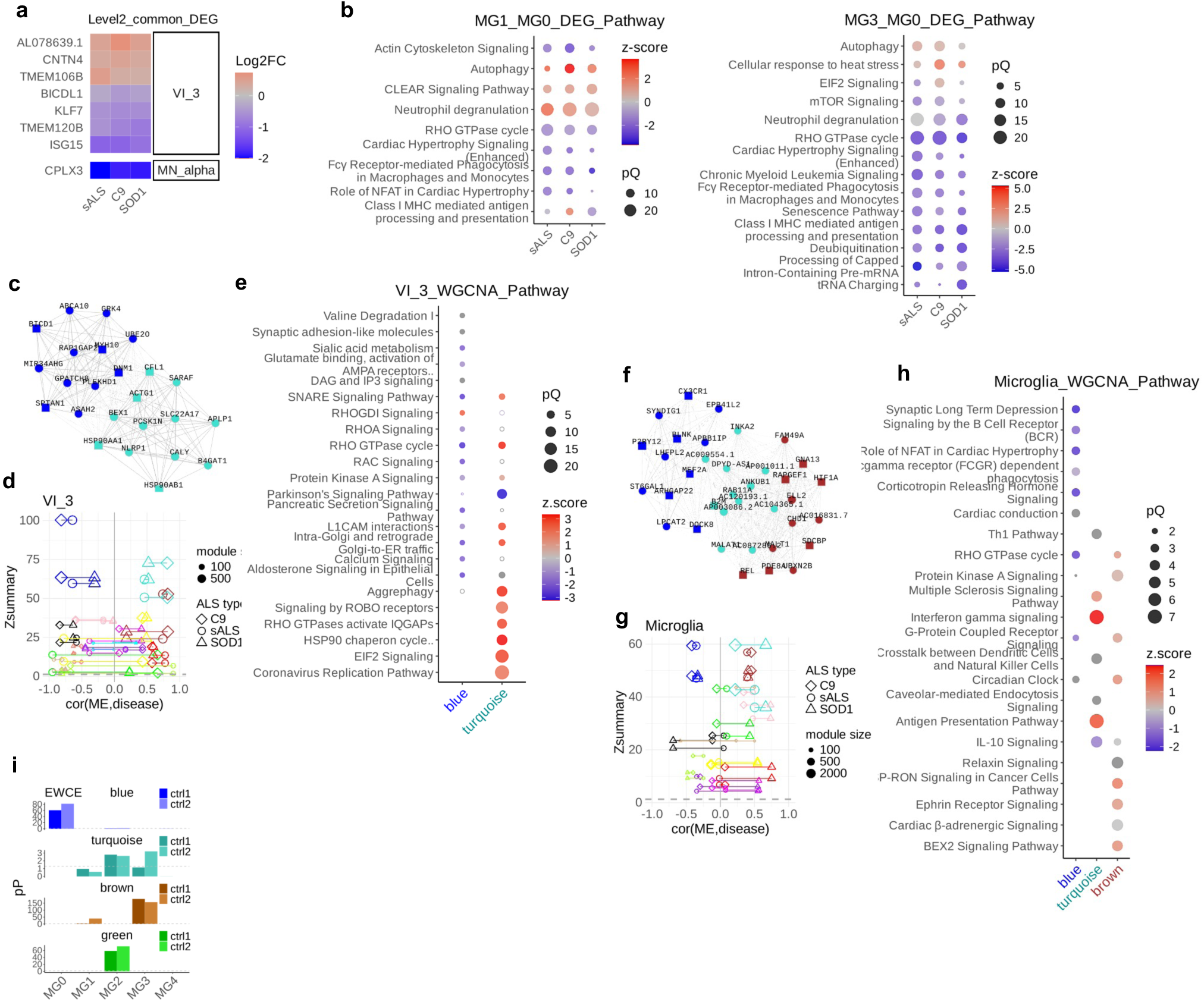
Common transcriptomic features across ALS types. (a) Common DEGs found in VI_3 and MN_alpha. (b) Top enriched pathways from DEG of (left) MG1 from ALS compared to MG0 from control (MG1_MG0), and (right) MG3 from ALS to MG0 from control (MG3_MG0), redundancy mitigated (Method), for each ALS type but plotted together. Note that we did not prioritize overlapping pathways. Meaning of pQ and z-score are the same as in Fig. 1. (c, f) Co-expression network diagram of top hub genes from hdWGCNA for VI_3 (level2) and microglia (level0), where square nodes represent genes that are part of the enriched pathways. (d, g) Preservation statistics (Zsummary) and the correlation between the module and disease status (cor(ME,disease)) for VI_3 and microglia, for all pairs of ALS types. Higher Zsummary means more similarity between modules, and larger absolute cor(ME,disease) means stronger association of the module to ALS. (e, h) Top enriched pathways for selected modules of VI_3 and microglia from hdWGCNA. As in (b), we did not prioritize overlapping pathways. Filled and open circles correspond to Q>0.05 and <0.05, respectively. (i) Enrichment of expression of genes from modules in microglia hdWGCNA, using EWCE.

Among microglia subtypes, MG1 and MG3, which carry neuroinflammatory and stress signatures, were increased in numbers in sALS and C9, respectively (Fig. 2d). Hence, regaining balance among these cell types, either at transcriptional or at population level, might be beneficial to people living with ALS. Hence, we focused on comparing MG1 and MG3 from ALS to MG0 from controls. To avoid the “double-dipping” problem, we used Countsplit^30^ to generate two statistically independent data sets from the snRNAseq data – “training” and “test” sets. The training set was used for re-labeling level2 types and the test set for DEG. MG2 was not detected, likely due to its scarcity. Despite only ∼20% of DEGs were overlapping across all ALS types (Fig. S18), there was strong agreement in the enriched pathways from MG1 and MG3 compared to MG0 across all ALS types (Fig. 4b). In the MG1-MG0 comparison, CLEAR signaling and autophagy was activated. Neurophil Degranulation shows opposing directions between the MG1-MG0 and the MG3-MG0 comparisons, suggesting this pathway was specifically activated in MG1. In MG3-MG0, the heat stress pathway was activated, which is the hallmark of MG3 (Fig. S9). These data shows that there are converging transcriptomics characteristics of microglia subtypes when compared to the homeostatic state across ALS types.

To further investigate preserved biological functions across ALS types, hdWGCNA^31^ was applied to the snRNAseq data (Fig. 4c-g). With the snRNAseq data, co-expression networks were constructed for each ALS type paired with its control, using the training set from Countsplit. Next, the networks are combined to identify “consensus modules”, which are sets of genes that are well-connected to each other in the combined network (Method). Also, “module eigengenes” (ME), which captures overall direction of expression of genes in a module, were calculated. The genes in the module are then applied to the test set to calculate module preservation scores (“Zsummary”^32^) across ALS types. The correlation between disease status and ME is also calcualted, and we focused on modules with high Zsummary and with consistent and strong ME correlation to ALS, across all ALS types.

In VI_3, blue and turquoise modules showed such characteristics, and they shared identical pathways enriched with opposing directions (Fig. 4d and e). SNARE Signaling is an example, where closer inspection reveals that genes in the inactivated and activated modules were related to synaptic vesicle release (SYT2/12, SNAP25, CPLX1, CAMK2G) and endocytosis (SYT4/11, CALM3, PRKACB, SNCG), respectively, indicating an imbalance between exo- and endocytosis. Intra-Golgi and Retrograde Golgi-to-ER Traffick is another example, where motor proteins were common to both modules. However, the inactivated module contains Golgi/endosomal genes (OGLIM4, GALNT2, MAN1C1, RAB30), while the activated module contains tubulins, highlighting involvement of different parts of the same pathway. In contrast, synaptic adhesion, calcium signaling, and heat-shock responses were specific to each module.

Enriched pathways in microglia modules showed different patterns. Genes in the blue, green, and brown modules were highly expressed in MG0, MG2, and MG3, respectively, while the those in the turquoise module was not high in a specific cell type (Fig. 4i). The green module was excluded from further analysis due to its weak correlation to ALS (Fig. 4g). It is not intuitive to understand the pathway enrichment results (Fig. 4h), but a few genes dominate most of the pathways. The enriched pathways in the brown module commonly include GNA12/13 and PDE4A/8A, indicating GPCR Signaling is the key driver. Most of well-connected genes in the turquoise module were lncRNAs (Fig. 4f), while enriched pathways commonly have up-regulated HLA-A/B/DRA/DRB1, indicating the “reactive” state of microglia^33^. The HLA region has genetic association (rs9275477) to ALS^18^ as well. In the blue module, the common genes were ITPR2, GUCY1A1, and PRKCD, with PRKCD being implicated in microglia from a Parkinson’s disease model^34^, though the role of the other two in microglia are not well understood. On the other hand, highly connected genes in the blue module were homeostatic microglia markers like CX3CR1, MEF2A, and P2RY12. Together, the co-expression comparison highlights inactivation of homeostatic microglial genes while the activation of MHC complex and GPCR pathways.

## Discussion

Previous ALS studies largely focused on motor neurons, microglia, and, to some extent, astrocytes. By comparing snRNAseq of sporadic, C9orf72, and SOD1-ALS, we show that the ventral inhibitory neurons undergo divergent transcriptomic changes in all three types of ALS, but also commonly exhibit impaired vesicle transport and synaptic function. The observations shed new light on the contributions of ventral neurons to ALS, which has been understudied. Identification of CPLX3 as a commonly downregulated gene in MN_alpha further demonstrates the importance of cell-type specific studies, which was undetectable at tissue level. Further investigation of cell type-specific gene expression in ALS in larger sample cohorts and incorporating improved motor neuron enrichment will be critical next steps toward fully dissecting the molecular changes in ALS.

## Data availability

All data supporting the findings of this study will be deposited in a publicly accessible repository (e.g., GEO) and will be made available upon publication. Processed data and analysis scripts will also be shared via an appropriate repository. Additional information is available from the corresponding author upon reasonable request.

## Author contributions

D.H. and W.C. designed the study.

D.H., S.C., G.Z. and C.L.W. performed snRNAseq data analysis

M.S., V.C., and Y.C. performed snRNAseq experiments.

D.H. and M.Z. performed RNAseq data analysis.

T.C. and R.C. performed RNAseq and snRNAseq library preparation and sequencing

A.J.G. and E.D.P. performed tissue preparation and pathology measurements

D.W. performed proteomics experiments.

D.W. and A.J.G. performed proteomics data analysis.

H.M. coordinated the project and genotyped samples

A.J.G. and H.M. acquired the tissues.

C.C. performed genotype analysis

J.H. and S-C.L provided biological consultations. D.H., S.C., and J.H. wrote the manuscript.

## Acknowledgement

This work was funded by Biogen. We thank the study participants and their family members. We thank Irina Leaf for coordinating sample purchase from University of Miami Brain Endowment Bank (funded by NIMH, NINDS, NICHD), Target ALS Postmortem Tissue Core for generously providing C9 and SOD1 samples, and Alexander McCampbell for discussions.

## Supplementary Figures

**Fig. S1.**
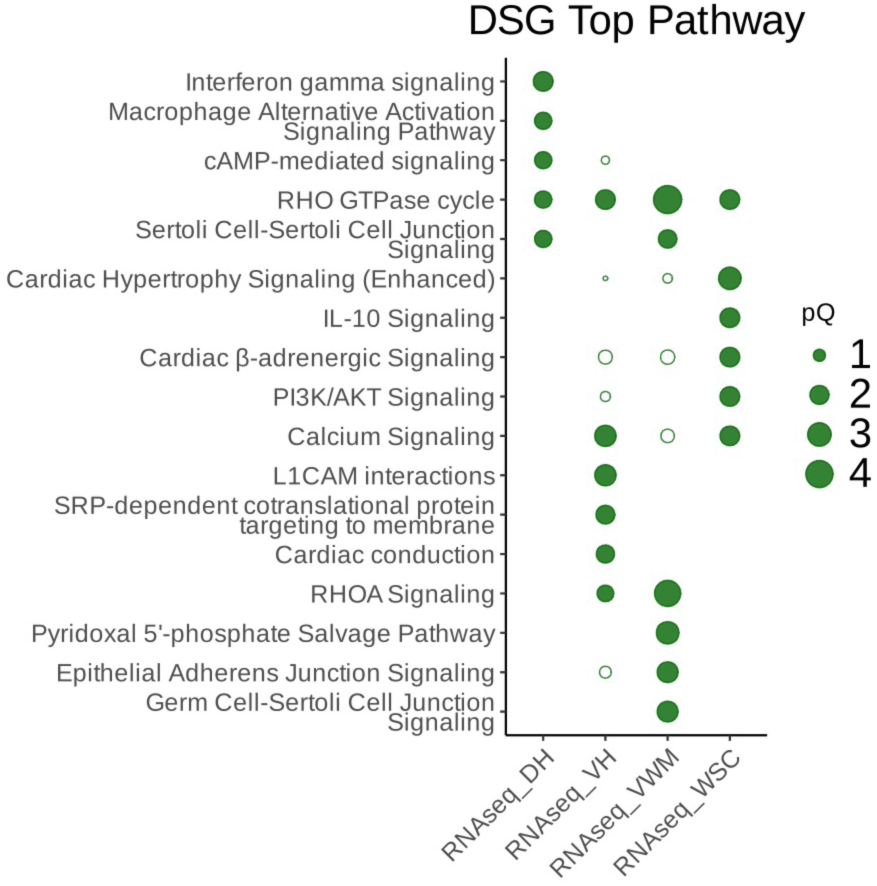
Top 5 enriched pathways from DSG per tissue from sALS RNAseq via IPA.

**Fig. S2.**
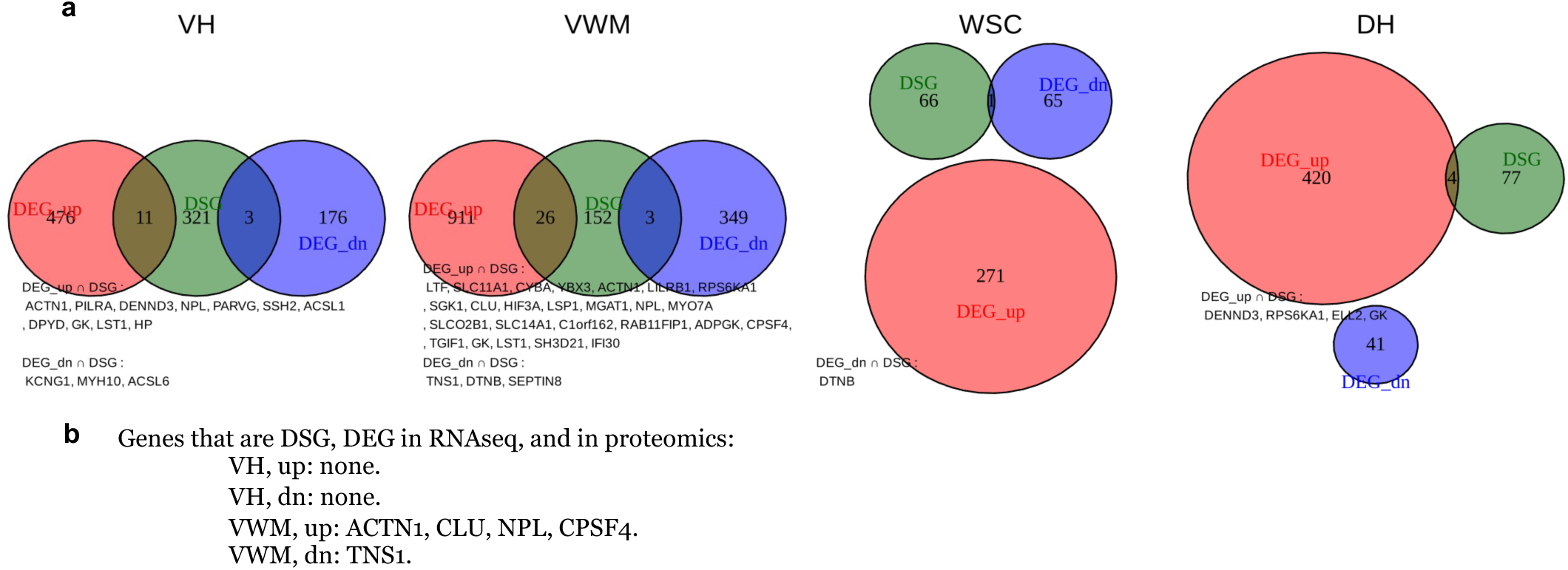
Overlapping DEG and DSG from RNAseq and proteomics. (a) Overlapping genes between DSG and DEG from RNAseq. (b) List of overlapping genes in DSG, DEG in both RNAseq and proteomics. Interestingly, four genes in VWM-up are all specifically expressed in astrocytes (data not shown).

**Fig. S3.**
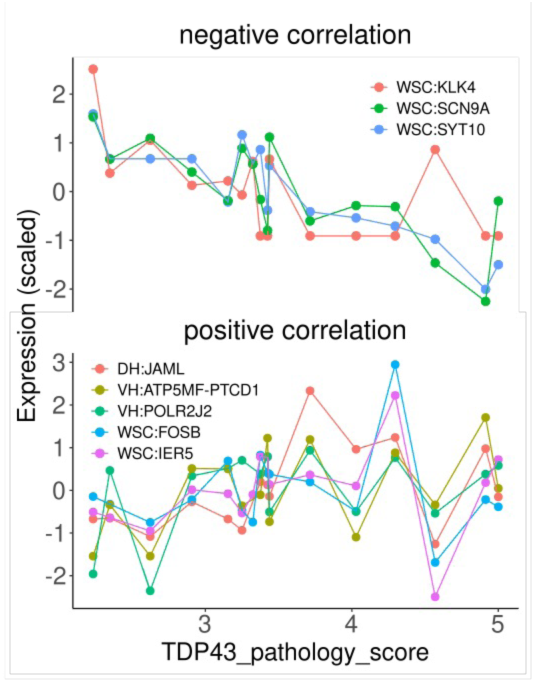
Gene expression correlated with TDP43 pathology in sALS at tissue level. In sALS tissues, RNAseq and proteomics data were regressed against TDP43 pathology using DESeq2 (Method). Genes with Q<0.05 and |Log2FC|>1.2 were filtered, and only 8 genes shown here passed it, all of which are from RNAseq but none from proteomics.

**Fig. S4.**
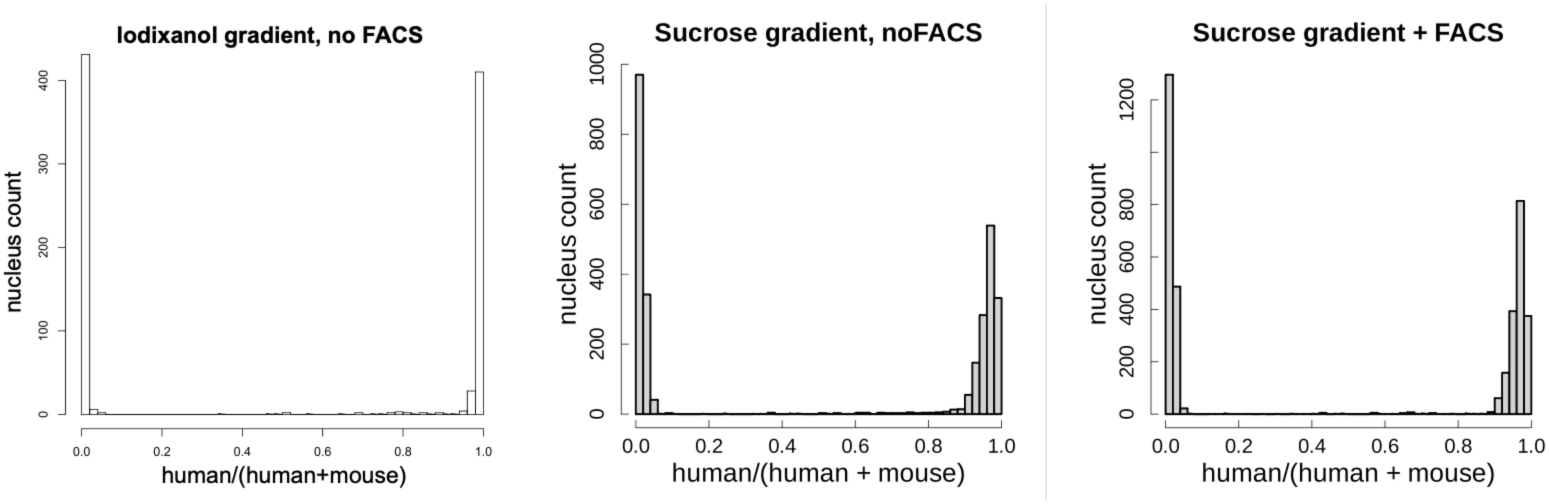
Species-mixing experiment for testing nucleus isolation methods. Nuclei from human and mouse spinal cords are isolated using the iodixanol gradient method, and sucrose gradient method with and without FACS. The isolated nuclei were subjected to snRNAseq, the reads were mapped to human and mouse reference genome simultaneously, and only uniquely mapped reads were retained. For each barcode that supposed to mark a single nucleus, the fraction of mapped reads to human genome is calculated (“human/(human+mouse)”). A histogram of this value is generated as in Klein et al, where thin bimodal plot without any spread is desired, which represents minimal cross-contamination among droplets. Our data show that the iodixanol gradient meets this criterion.

**Fig. S5.**
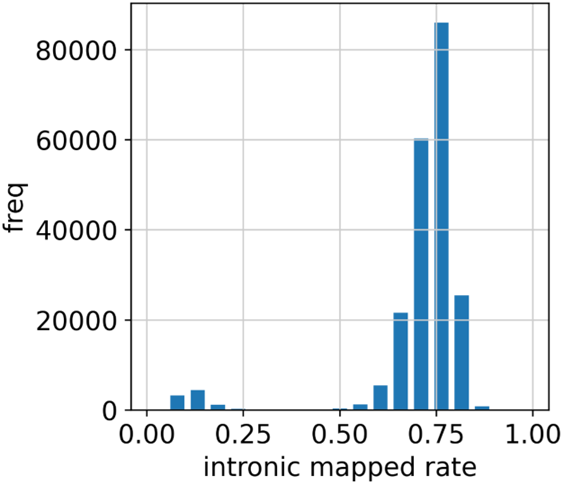
Histogram of intronic mapped read rate. <0.4 are considered originating not from nuclei, hence removed.

**Fig. S6.**
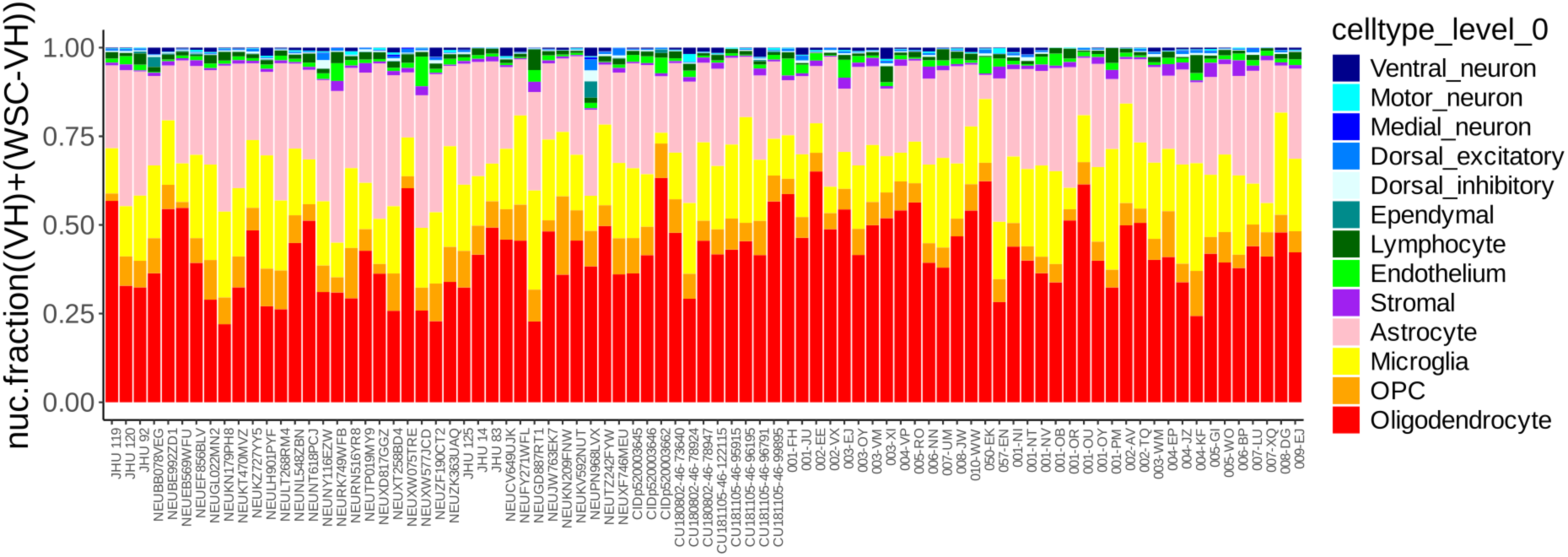
Cell type proportion for each sample in snRNAseq, nuclei from VH and WSC-VH per sample combined.

**Fig. S7.**
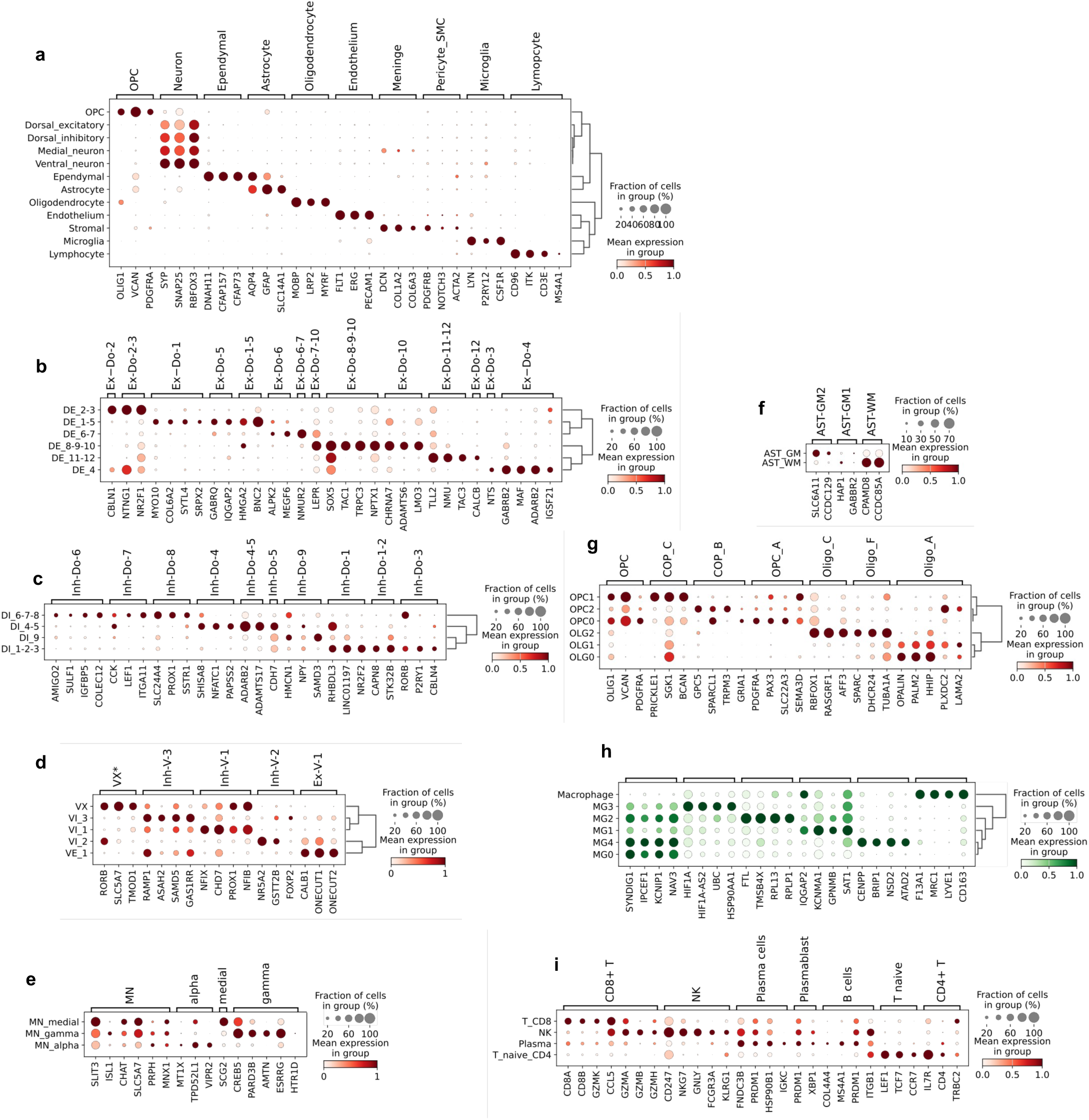
Marker gene expression in the clusters. (a) Level0 markers. (b-g, i) Level2 markers. (b, c, d, f): Dorsal excitatory (DE), dorsal inhibitory (DI), ventral neuron (V), and astrocyte (AST) level2 markers. In (d), VX* is not listed in the reference, hence 1-vs-rest t-test is used to identify the markers. (e) Motor neuron level2 marker. (g) Oligodendrocyte (OLG) and OPC markers (h) Level2 for microglia are defined by de-novo clustered combined with marker gene identification (green dots) and GO enrichment (Fig.S9) (Method). (i) Lymphocyte level2, with marker genes from a variety of sources.

**Fig. S8.**
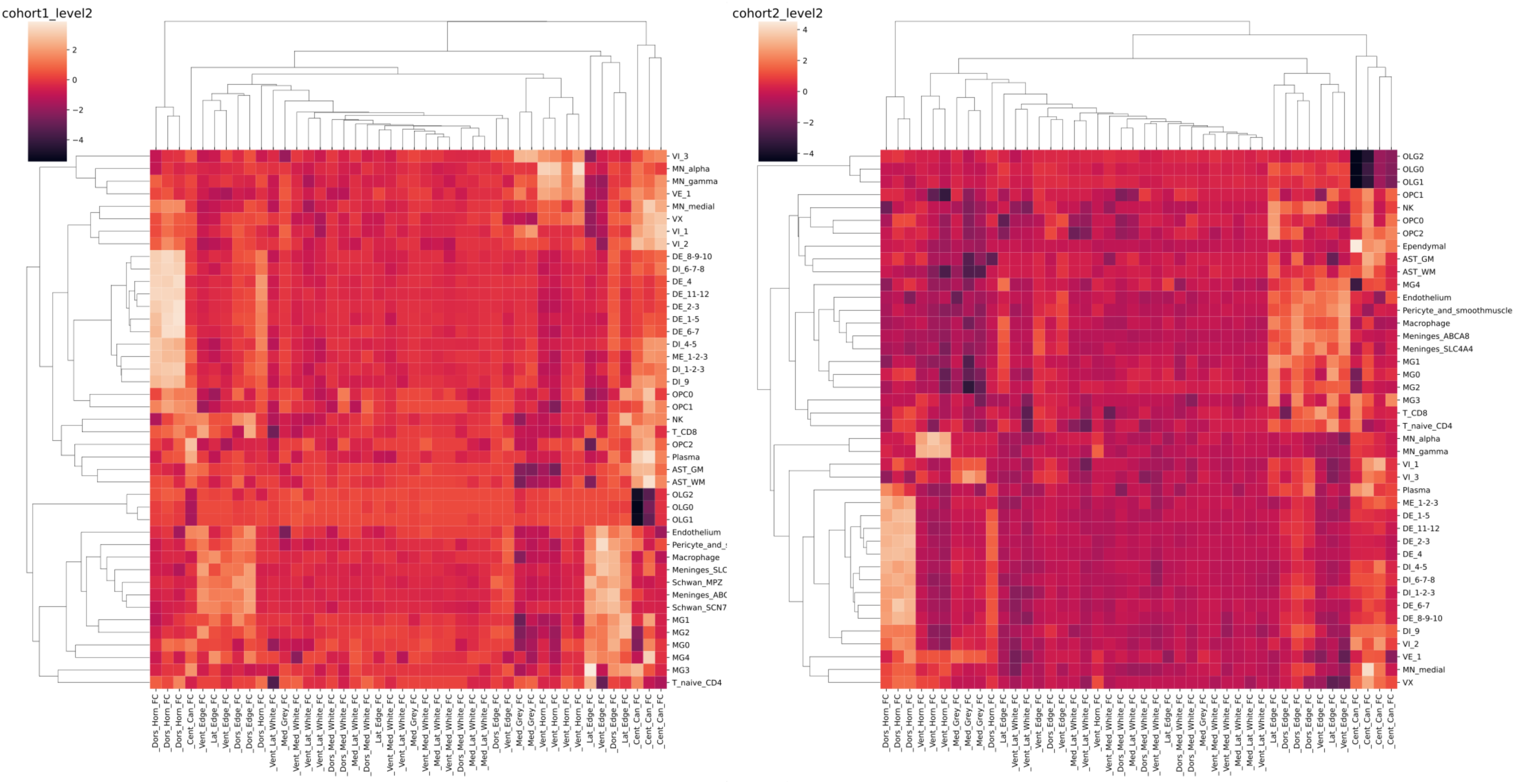
Gene expression correlation of cohort1 and 2 controls and spatial transcriptomics (ST) data. Top 90 genes with high specificity scores with average UMI>0.5 in the cohort1 and the cohort2 control groups are correlated with gene with BF>0.8 in the ST data per region.

**Fig. S9.**
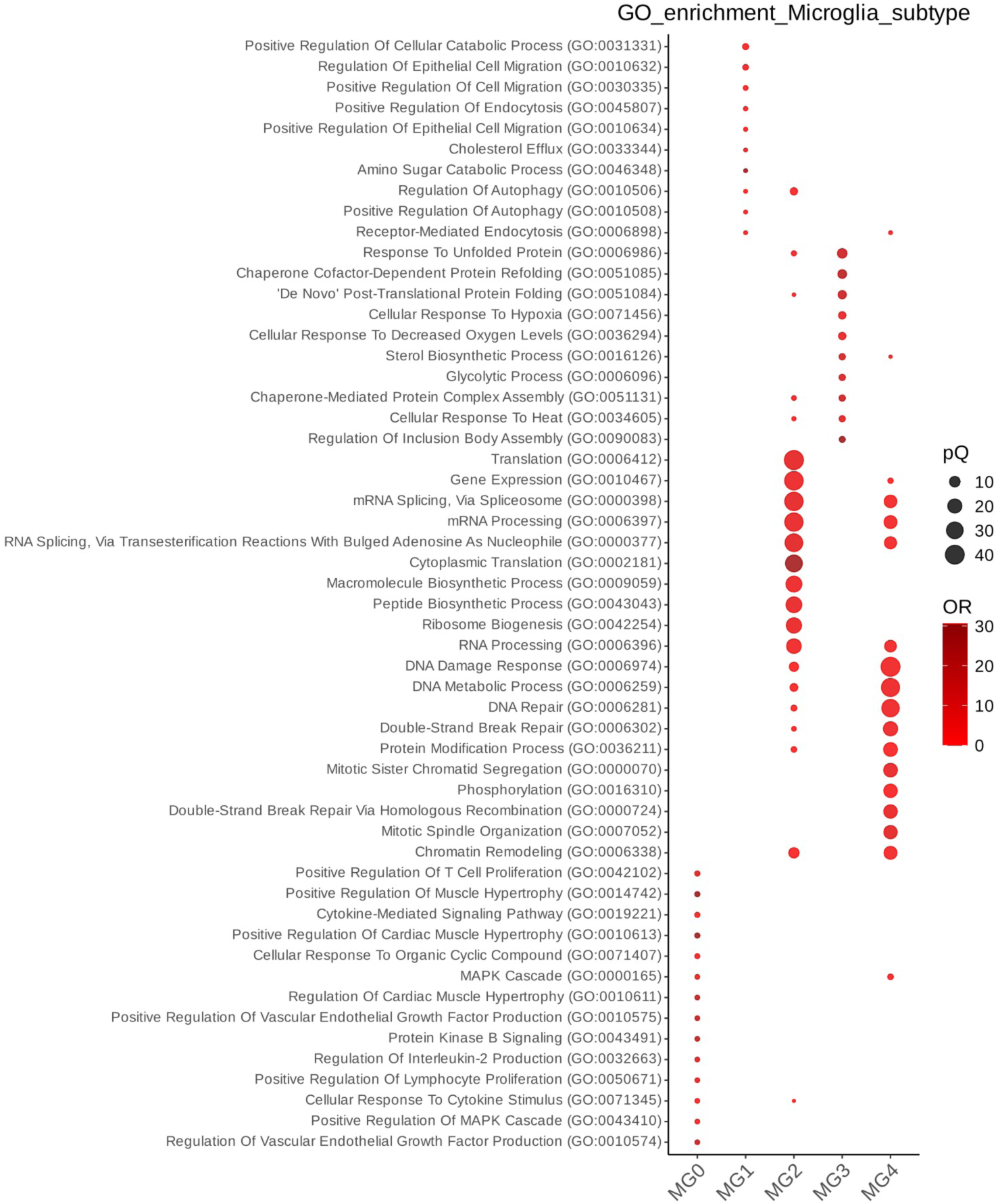
GO term enrichment in microglia level2 types. Among microglia level2 types, 1 vs rest Wilcoxon rank test is performed, and genes with Q<0.05 and |Log2FC|>1.5 are subjected to GO enrichment analysis using EnrichR. Top 10 pathways with smallest Q values for each cell types are selected, and plotted together.

**Fig. S10.**
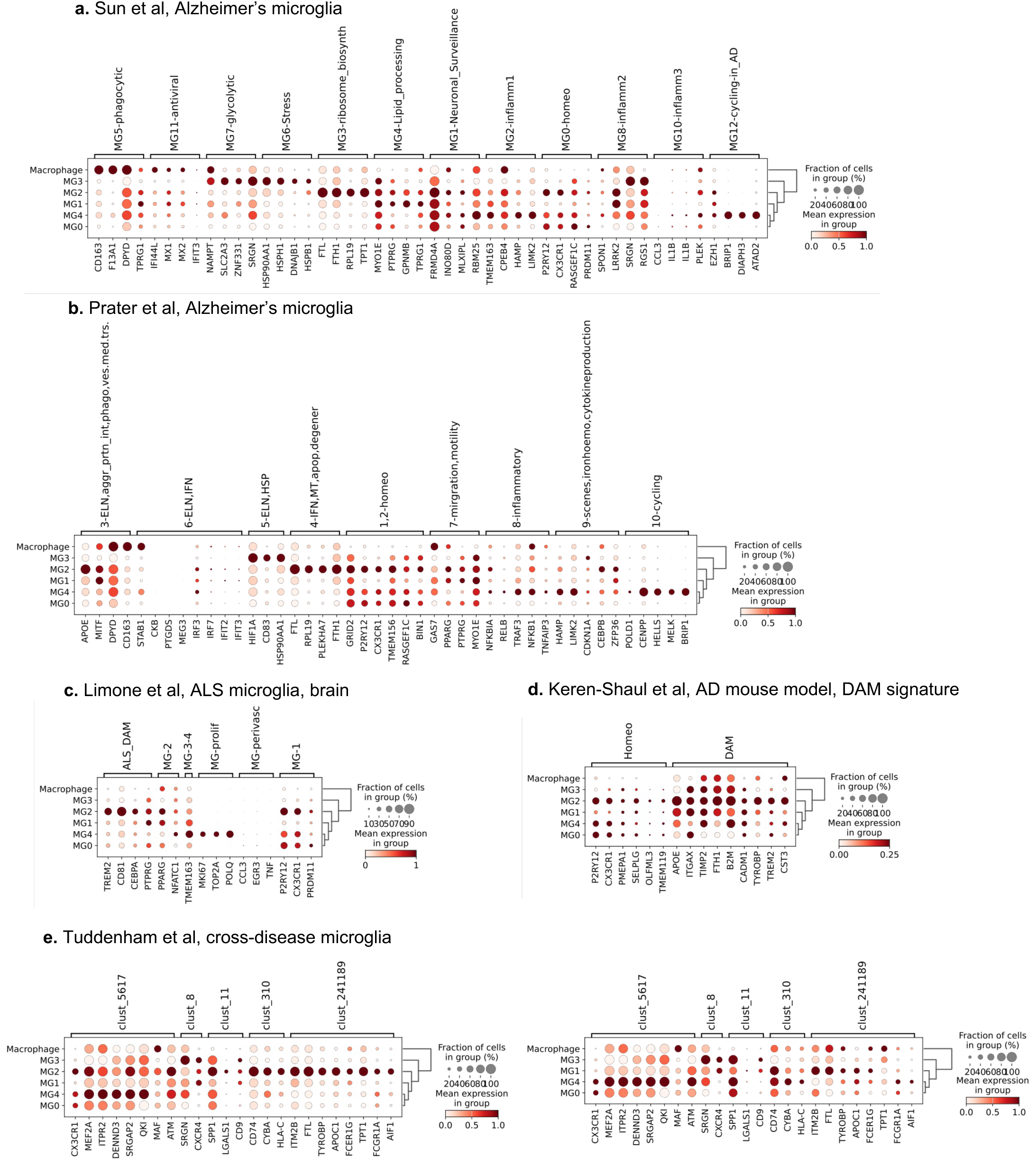
Expression of marker genes from literature in the microglia level2 clusters. (a, b) Marker genes of microglia from Alzheimer’s brain cortex. (c) from ALS brain cortex, (d) from AD mouse model that defines the disease-associated microglia (DAM) signature, which is not confined to a single cluster. (e) Marker genes from a cross-disease study of microglia, with and without MG2.

**Fig. S11.**
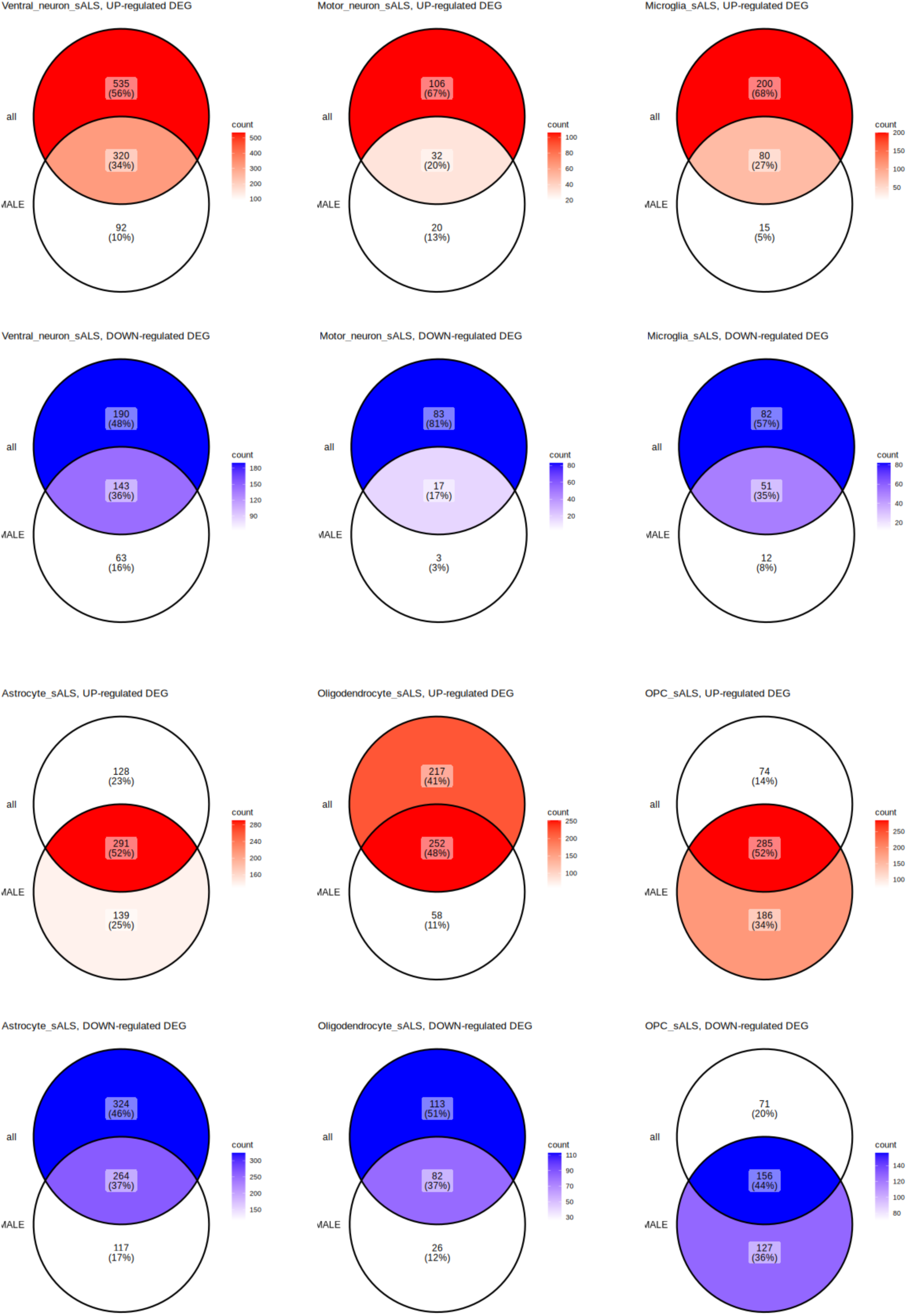
Comparison of DEG from all ALS vs control and male ALS vs control, for selected cell types in sALS.

**Fig. S12.**
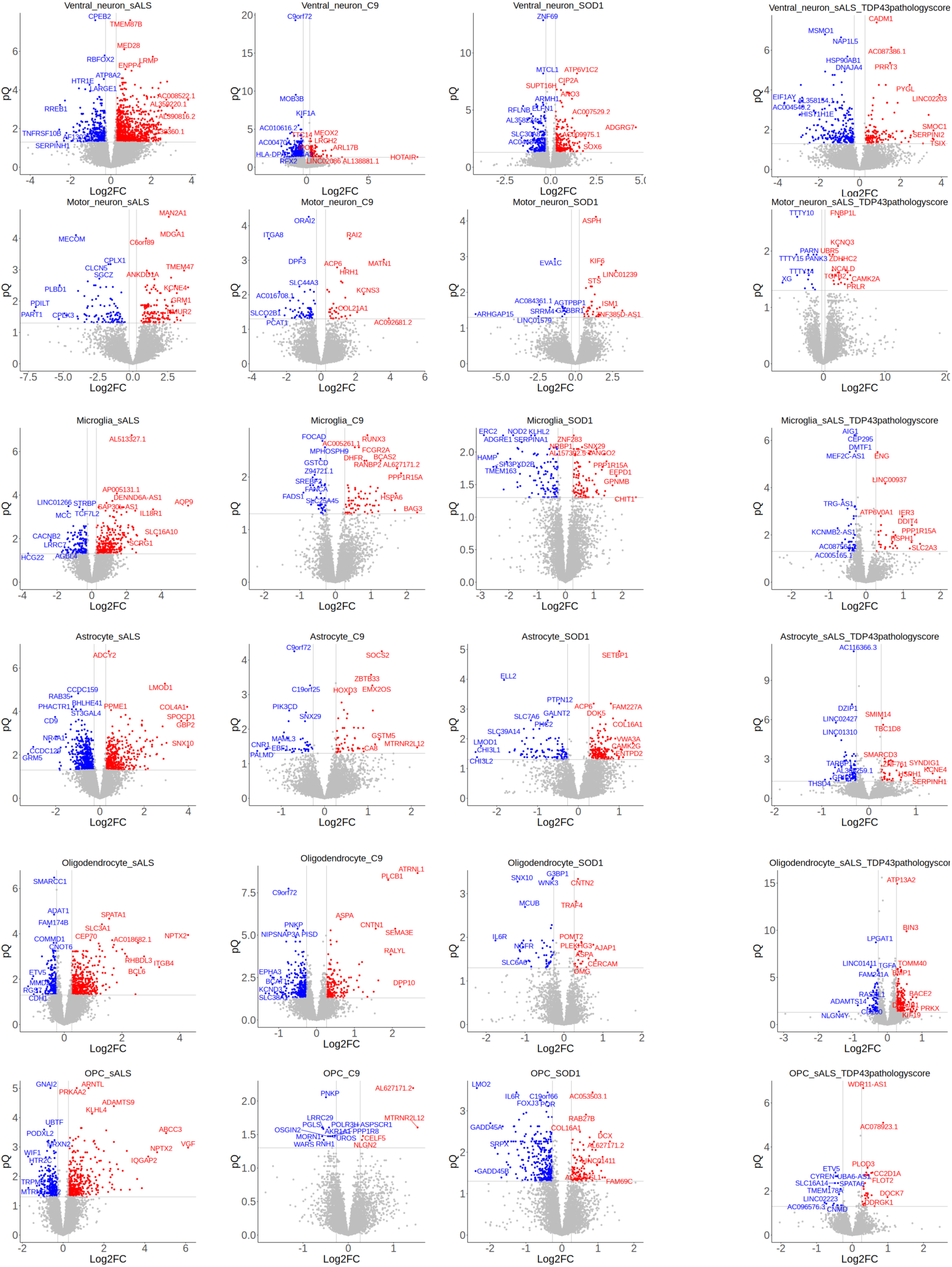
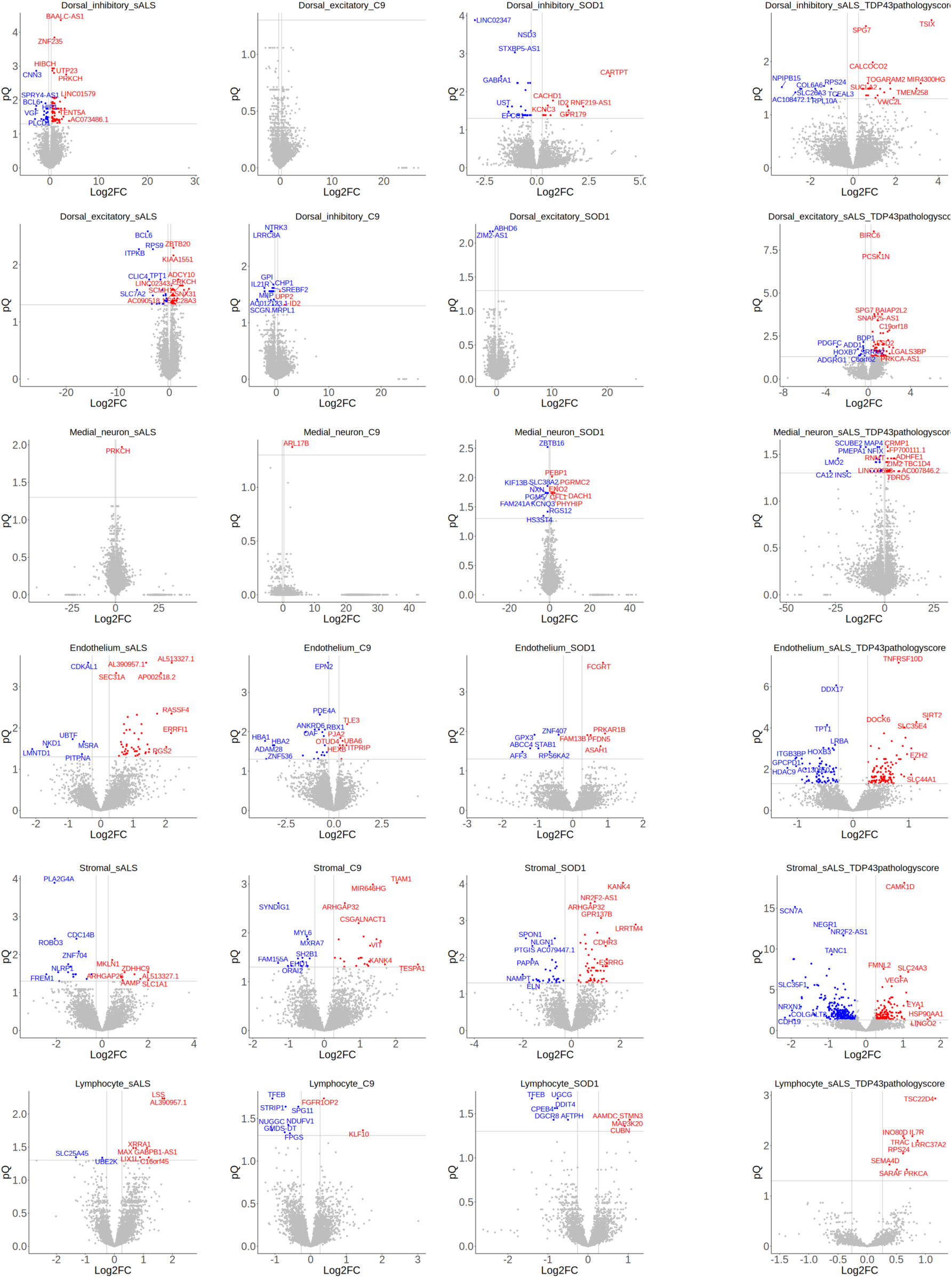
Volcano plot of snRNAseq per cell type, level0. Horizontal and vertical lines denoting pQ=0.05, |Log2FC|=Log2(1.2), respectively.

**Fig. S13.**
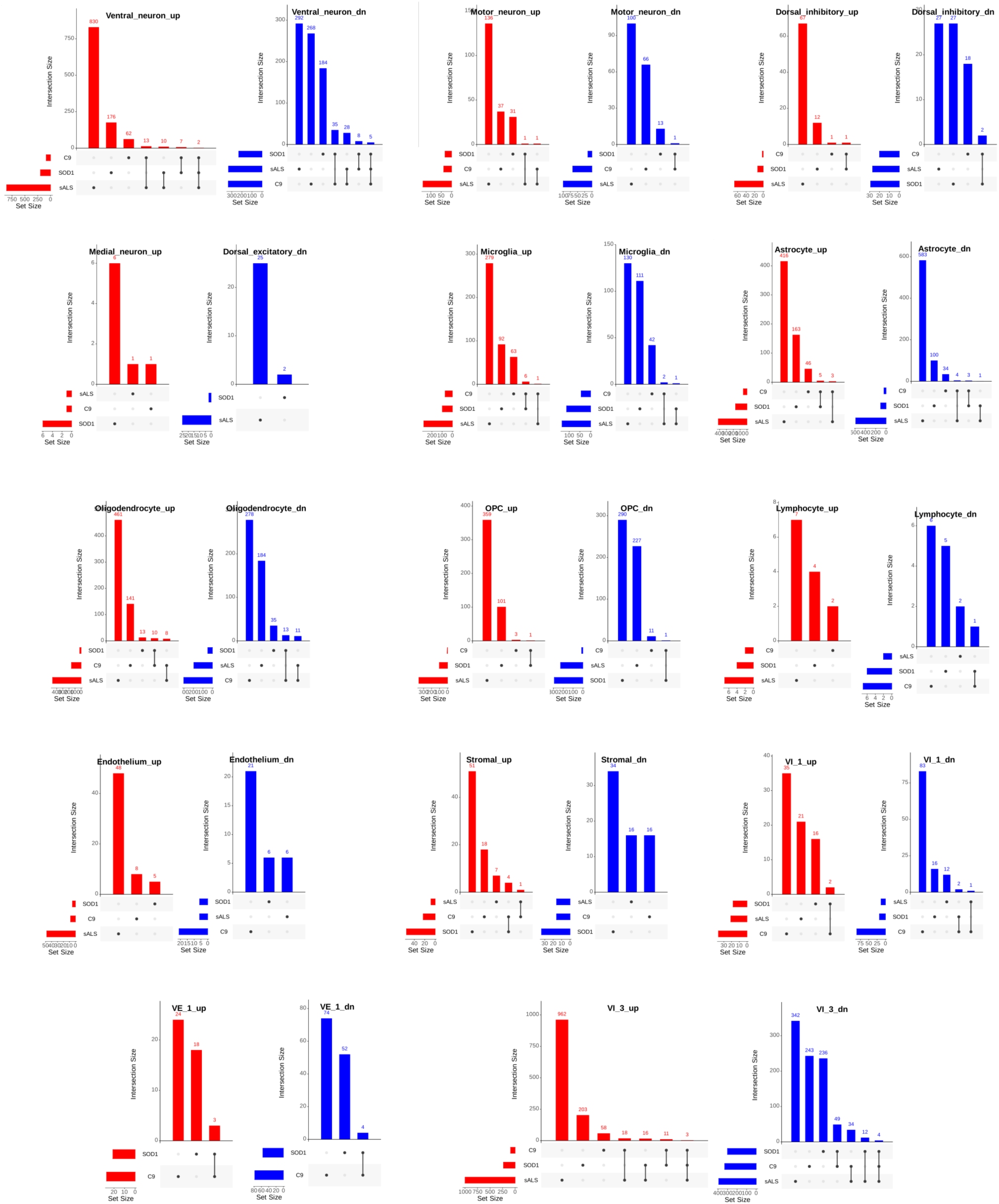
Upset plot of snRNAseq per cell type at level0. There are few common DEGs across all ALS types.

**Fig. S14.**
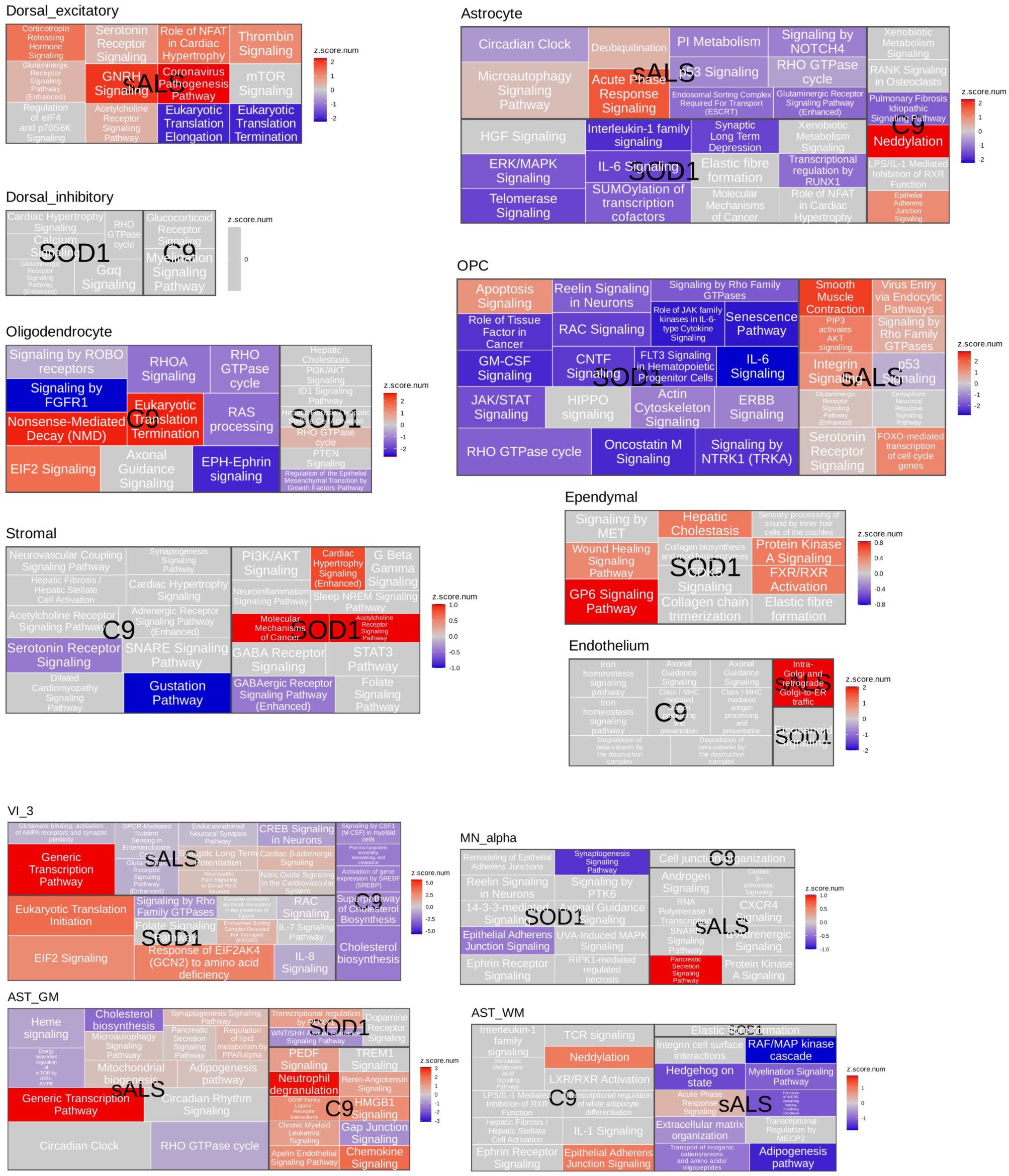
vPathway analysis of DEG from snRNAseq. Top enriched pathways from DEGs (IPA) for level0 and selected level2 cell types.

**Fig. S15.**
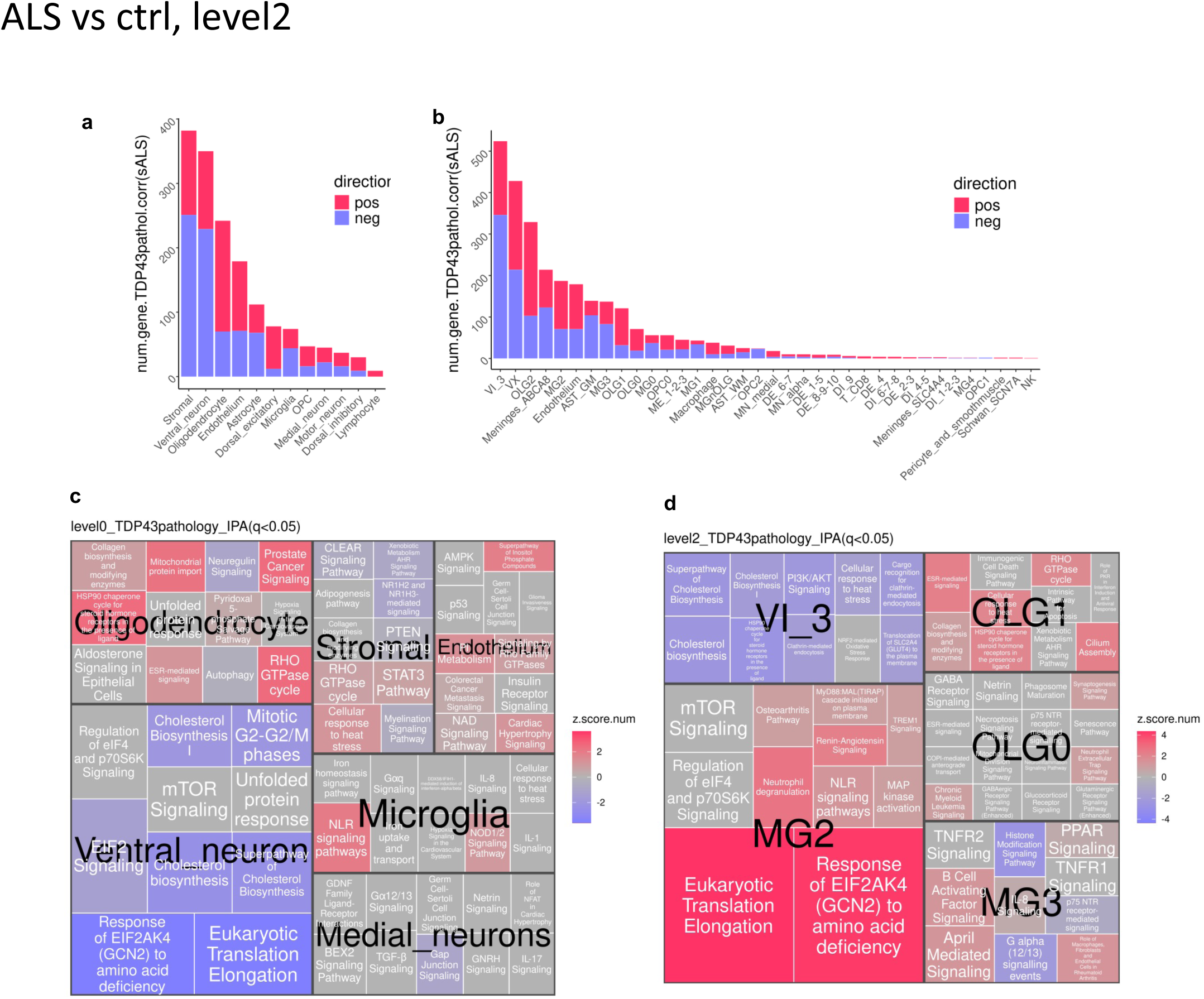
Number of genes that correlate with TDP-43 pathology (p.adj<0.05, |Log2FC|>1.2), and the corresponding top pathways (IPA) for level0 and level2 cell types.

**Fig. S16.**
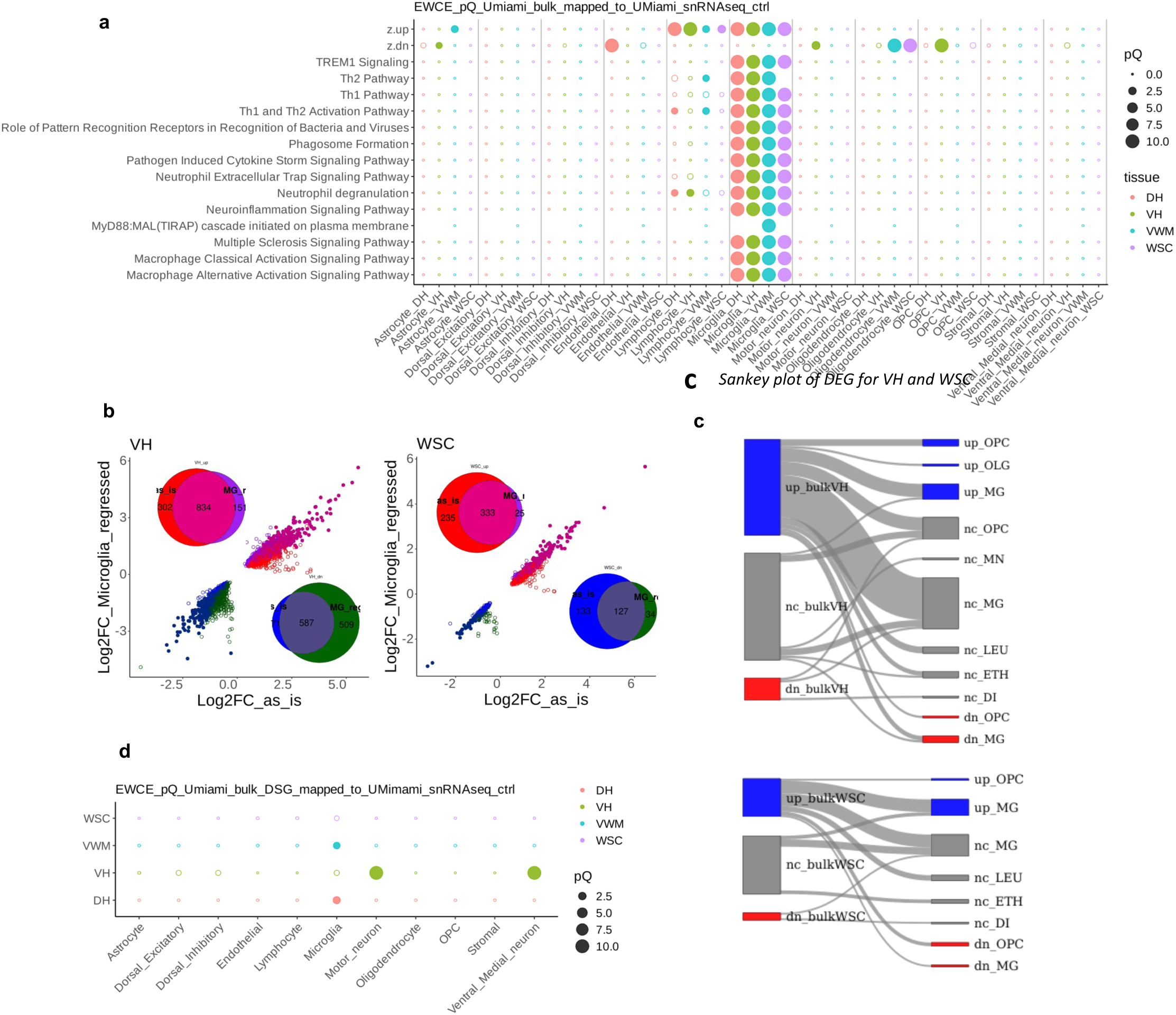
Reanalysis of sALS RNAseq DEG and pathways with snRNAseq data. (a) EWCE of genes from DEG and top 10 pathways, using expression level of control samples from cohort1. (b) Log2FC plots of DEG with and without microglia fraction regression. Inset: Venn daigram up- and down-regulated genes in each tissue. (c) Shanky plot of enriched pathway between sALS RNAseq and snRNAseq. “up” and “dn” correspond to z-score>1 and z-score < -1, and the rest are “nc”. (d) EWCE of DSG using expression level of control samples from cohort1. In all the plots, filled and open circles represent Q>0.05 and <0.05, respectively.

**Fig. S17.**
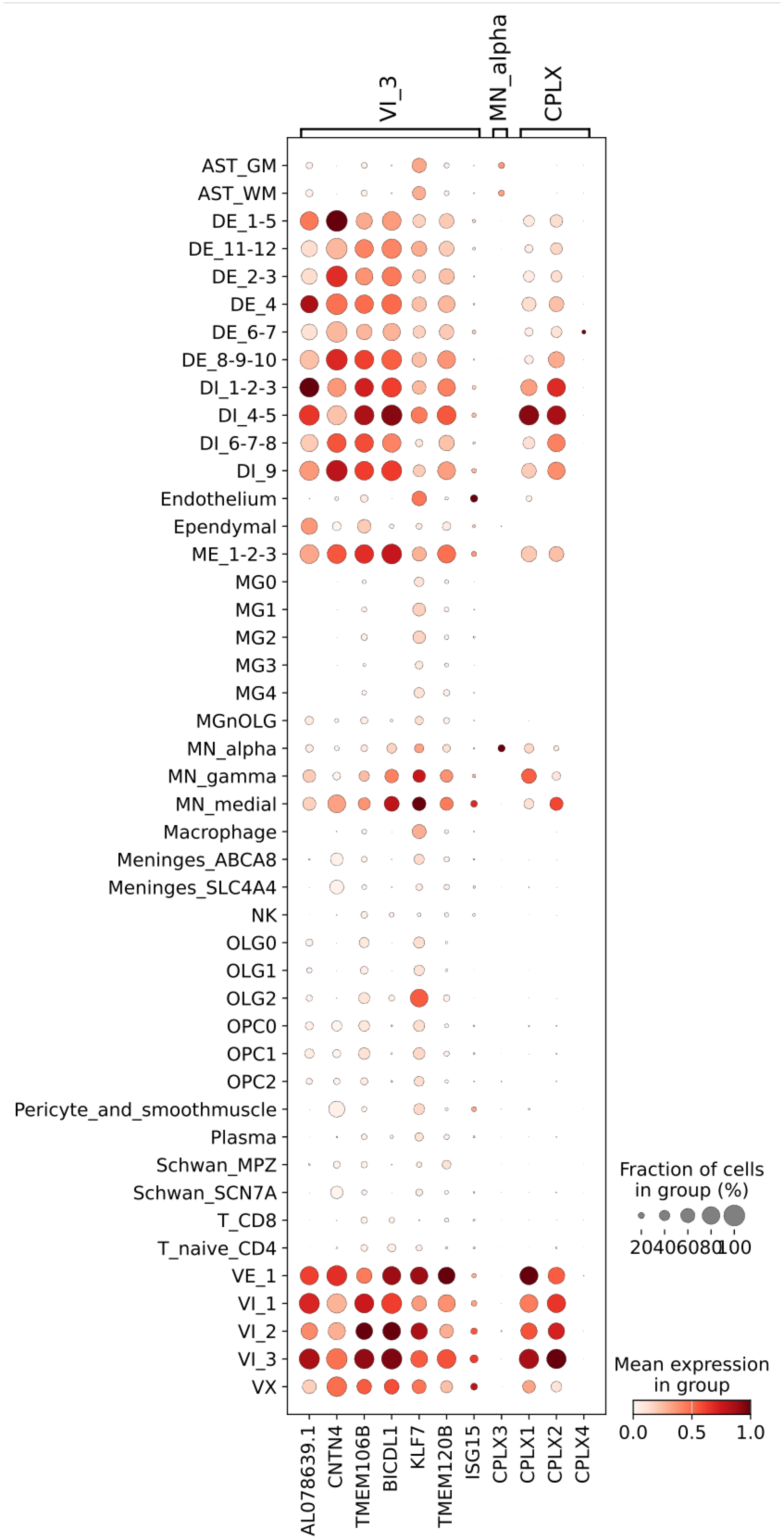
Expression level of CPLX3 and other genes for comparison. Genes in “VI_3” are common DEGs of VI_3. Those in “CPLX” are CPLX1, CPLX2, and CPLX4 to be compared with CPLX3. “MN_alpha” categroy is used for CPLX3.

**Fig. S18.**
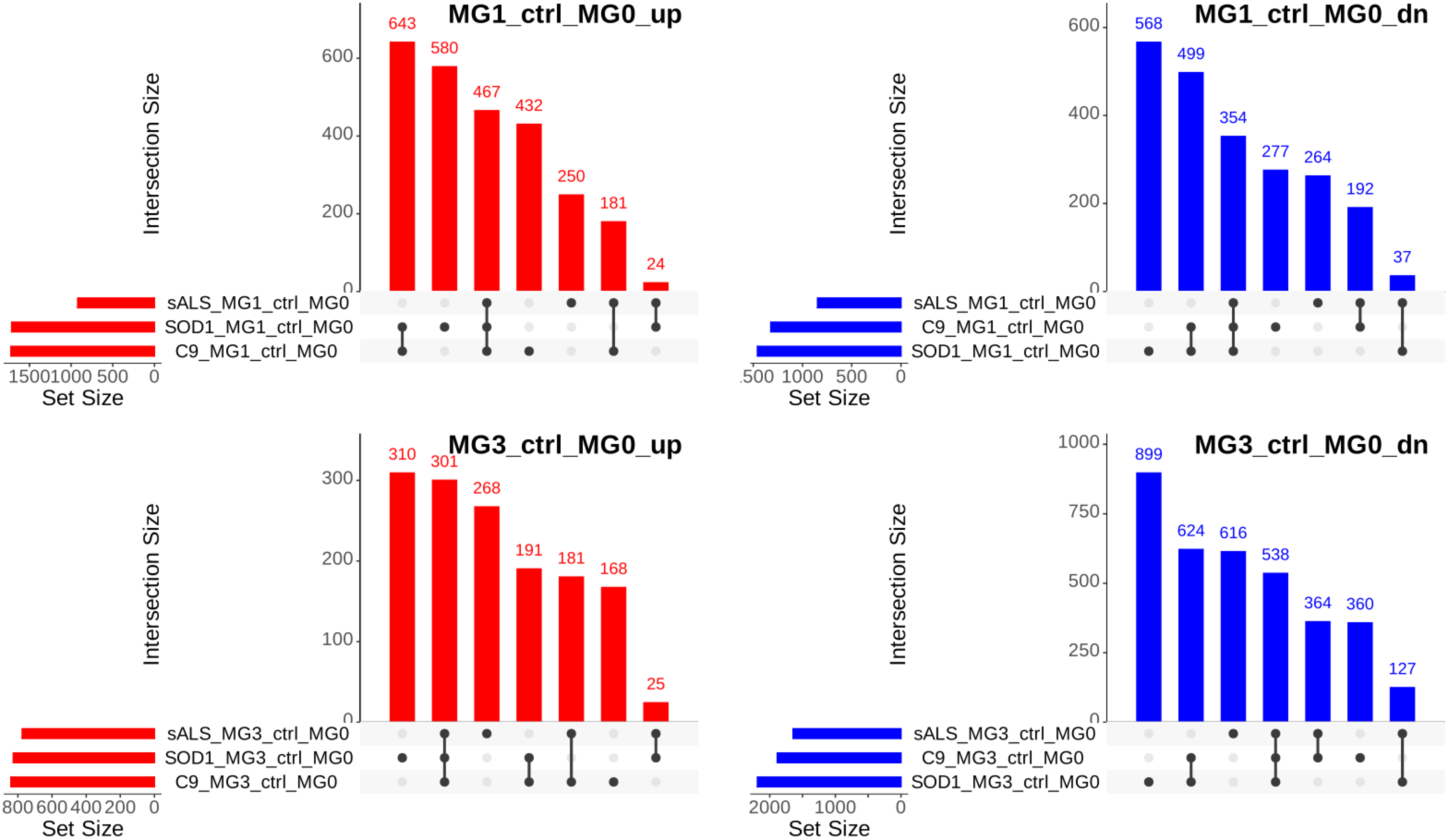
Upset plot of DEG in ALS MG1 and ALS MG3 compared to control MG0.

**Fig. S19.**
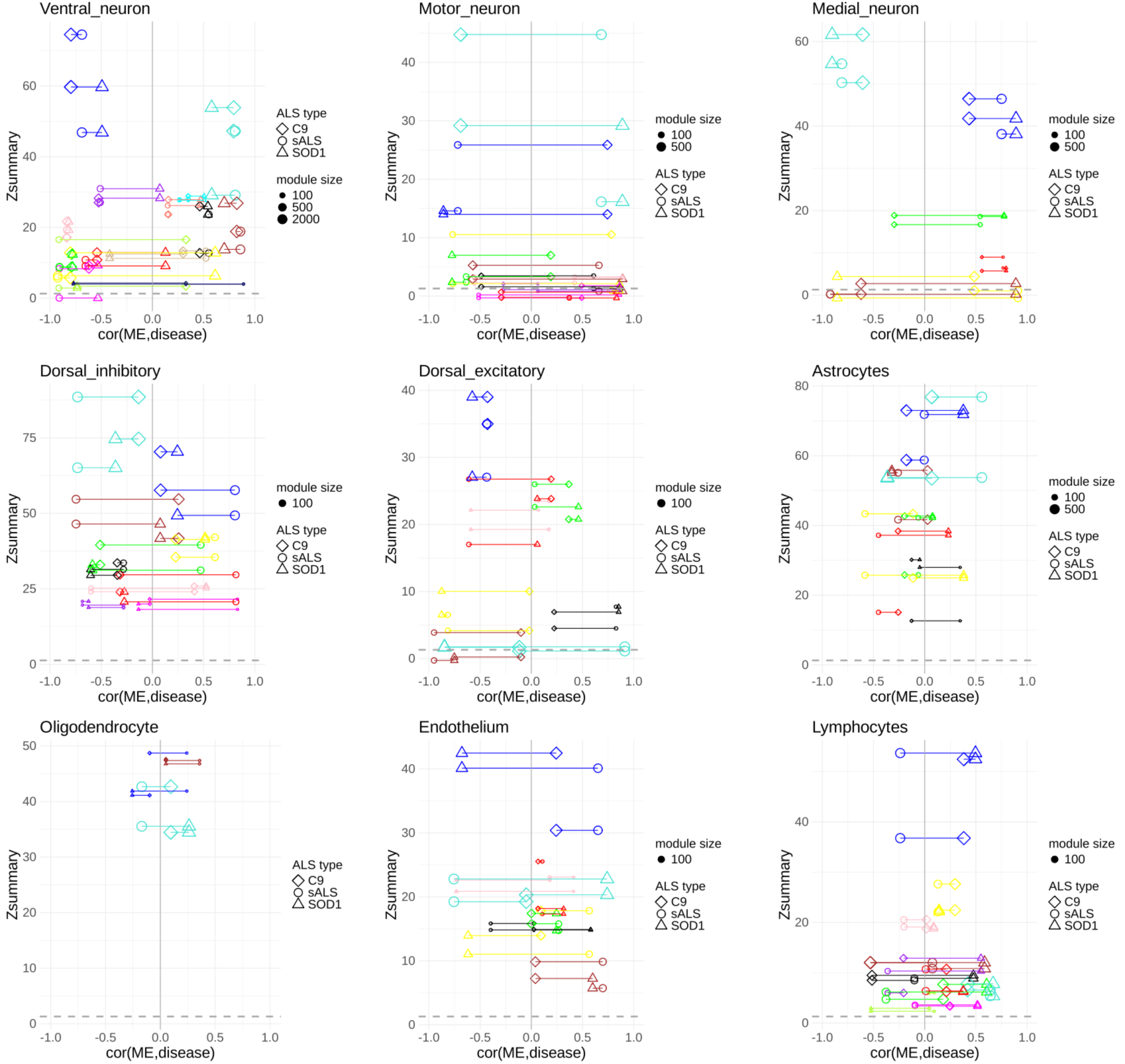
Preservation statistics (Zsummary) and the correlation between the module and disease status (cor(ME,disease)).

