## Supplementary material for "Comparative snRNAseq study of C9orf72, SOD1, and sALS spinal cord": Method

**Spinal cord tissue processing** For the sALS, frozen transverse sections of lumbar spinal cords from sALS and non-neurological control subjects were purchased from the University of Miami Brain Bank. Hexanucleotide repeat expansion of C9ORF72 was measured (Asuragen AmplideX® PCR/CE C9orf72), and two samples – 012CZ and 004HI - had the repeats, hence were excluded from downstream analysis. Other genetic variants association with familial ALS were investigated using targeted sequencing, and no such variants were found (cohort1_sample_description.tsv, "genotype-" columns). For RNAseq, VH, DH, and VWM were isolated using 1mm-diameter tissue punch for each sample with 1.5mm thickness, and adjacent whole section (WSC) was also prepared for further processing. Meninges were removed as much as possible using a razor blade prior to punch and sectioning. For proteomics, VH and VWM were punched from adjacent sections of the tissues, with same dimension. For snRNAseq, VH were punched (1 mm diameter x 3 mm depth), yielding two VH tissues per sample, and the remainder (WSC-VH) were sectioned into 1mm thickness, all of which were subjected to nucleus isolation. Frozen lumbar spinal cords from C9 and SOD1 were purchased from Target ALS consortium, and controls samples from Cureline and NBS (cohort2_sample_description.tsv). Tissue processing was similar to that in sALS, except that the VH punch was 1 mm diameter x 1.5 mm depth x 2 rounds. Genotype information was obtained from Target ALS consortium, and no further genotyping were performed.

**RNAseq sample preparation** RNA was isolated using the QIAgen miRNeasy Micro Kit (QIAgen # 217084) using QIAzol-based lysis, and on-column DNase treatment according to manufacturer's instructions.Libraries were prepared from 100 ng of RNA using the KAPA RNA HyperPrep Kit with RiboErase (HMR) (Roche, #KK8561) according to the manufacturer's instructions. The following parameters were used: RNA fragmentation at 94°C for 3 min, adapter ligation using KAPA UDI Indexes (Roche, #08861919702) at 1.5 µM, and 14 cycles of PCR. Libraries were QCed on a LabChip GX using DNA HS Reagent Kit (Revvity, # CLS760672), normalized, and pooled for sequencing. Pools were sequenced on a HiSeq 2500 (Illumina), in paired-end mode 2x75bp, targeting 40 million fragments per sample.

**RNAseq Data Analysis** The reads were aligned to Gencode GRCh38.v34 reference using STAR[1] (v2.7.2d), and quantification via RSEM[2] (v1.3.2). Data quality metrics were generated with RNASeQC[3] (v1.1.8). Differential expression analysis between sALS and control samples was performed with DESeq2[4] (v1.34.0) with RIN, sex, and ethnicity included as covariates, implemented trough Expression Analysis[5]. Overdispersion was estimated using all the samples, and the function apeglm was used for LFC shrinkage to reduce noise and preserve large differences[6]. Differentially expressed genes (DEG) refers to genes with $|\log_2 FC|>1.2$ and adjusted p-value (Q) < 0.05, throughout the paper. Pathway enrichment analysis of DEG was performed using Ingenuity Pathway Analysis (IPA). Pathways with 2 or fewer genes were excluded. The Q values were calculated using the Benjamini-Hochberg method.

To identify differential local splice events between sALS and control tissue, we ran rMATs-turbo[7] (v4.1.2), Leafcutter[8] (v0.2.9) and Majiq[9] (v2.4), with Gencode GRCh38.v34 reference, and took the union of all the reported events. We define "FDReq" as adjusted p-values for rMATS and Leafcutter, while 1-P(dPSI) for Majiq for convenience. For Leafcutter, a cluster is dissolved into all possible three cassettes (triplet), and dPSI is recalculated according to Majiq's definition for each triplet, so that the result can be compared to the other two algorithms. For rMATS, we excluded events with maximum average junction counts per group is smaller than 50, as those might yield potential false-positives. For the identical events identified in multiple algorithms, we select one event that (1) has the maximum |dPSI| if FDReq<0.05 and (2) has minimum FDReq and the corresponding dPSI if FDReq>0.05 (manuscript in preparation). Note that "section" in the sample sheets and meta data is identical to WSC.

**Proteomics sample preparation** Frozen tissue punches were resuspended in 150µl of lysis buffer (4% SDS, 0.1M Tris/HCl pH 8.5, 5 mM EDTA ) and incubated for 5 min at 95°C in a Thermomixer. Samples were sonicated using a Covaris E220 for 20 cycles of 2 sec at maximum energy output, briefly heated, then centrifuged to remove insoluble material. Protein concentrations were determined by BCA. 20 µg of protein per sample were incubated in lysis buffer supplemented 1:10 with 0.1M dithiothreitol (DTT) for 10 min at 70°C, then alkylated at room temperature for 30 min with 0.3M iodoacetamide (IAA, added at 1:10). Digestion was carried out with modified trypsin (Promega, 2µg/sample) supplemented with 0.1µg LysC using the S-Trap tips packed with Empore C18 extraction discs (CDS, Cat. No. 98060402173EA) and Glass Microfiber Filter discs (Whatman, Article No. 28418314). Eluted peptides were dried and resuspended in 0.1M triethylammonium bicarbonate (TEAB).

**Isobaric peptide labeling with Tandem Mass Tags (TMT)** 10% of peptides were pooled to generate bridge channels for either VH or VWM to be included in each is Tandem Mass Tage (TMT)-labeled set. The remaining peptides from each sample (~90% of all peptides) were individually labeled with 0.2mg TMT 11-plex reagents (ThermoFisher Scientific, Lot No.: UJ292056) according to the manufacturer's recommendations such that samples from both ALS and CTL subjects were equivalently distributed across channels within a given 11-plex TMT set.

**Peptide fractionation and LC-MS/MS analysis** Samples were resuspended in 440µl 0.5% trifluoroacetic acid (TFA), sonicated, clarified, fractionated by high-pH reverse-phase chromatography on an Agilent 1290 Infinity LC system. Fractions were combined using a non-continuous pooling scheme, resulting in 24 fractions. Fractions were dried and desalted using C18 StageTips. Peptides were resuspended in 22ul 0.1% TFA. 4µl per fraction were injected and analyzed across a 140min nLC-MS gradient on a QExative HF mass spectrometer (ThermoFisher Scientific) using a fixed injection time method with a resolution of 60,000 for MS2 scans. Resulting MS/MS data were processed in MaxQuant and searched in Andromeda against a comprehensive UniProt/SwissProt human protein database (Download date: 30 Sept 2019).

**Proteomics data analysis:** After QC and median-normalization, Limma was used for DEG analysis, using sex, age, and RIN as covariate for VH and VWM, separately. Note that "VGM" in the sample sheets and meta data is identical to VH. Pathway analysis was done as in RNAseq Data Analysis

**TDP43 pathology: tissue processing, staining, and imaging**: FFPE blocks were prepared from ~3mm-thick transverse spinal cord blocks were drop-fixed in 10% NBF for 48hr followed by standard dehydration and paraffin infiltration. Blocks were gently faced and then sectioned at 5um sections for TDP43 staining. Immunohistochemical detection of TDP43 was carried out using a mouse monoclonal anti-TDP43 antibody (Biogen, L95A-42 TDP43-unconjugated) at a concentration of 0.25µg/ml as the primary antibody and an OMAP anti-mouse HRP as the secondary antibody for detection. Ventana Discovery Purple was used as the chromogen for visualization. Images were acquired on a 3DHistech Pannoramic-250 full slide scanner at 20x.

VH regions of interest (ROI) were manually annotated in Visiopharm for each tissue sample. **Total TDP43 protein abundance in ventral horns:** TDP43 area was detected using custom apps to detect TDP43 signal and corresponding tissue area. TDP43 expression per donor was determined based on relative TDP43+ area within the ventral horn ROI. **TDP43 inclusion frequency:** TDP43 inclusion frequencies were determined by manual inspection of individual MN cell crops from each donor and are reported as the % of MNs with inclusions present relative to the total number of MNs sampled per donor. **Semiquantitative TDP43 pathology scoring:** Individual MN cell crops were classified into groups based on qualitative assessment of degree of TDP43 pathology and assigned a numeric score: normal appearing (no apparent pathology) = 1; mild pathology = 3; moderate pathology = 5; and severe pathology = 7. For per donor metrics, the mean numeric score across all sampled MNs (~50 per sample) is reported.

**snRNAseq sample preparation** Nuclei were isolated using a modified version of methods outlined by Hrvatin *et al*[10]. VH and the remaining tissues from human spinal cord section were briefly thawed on ice, homogenized in chilled 0.1% dithiothreitol, 0.001% Triton-X 100, 0.125% RNAse inhibitor in Tris-HCl. Homogenized human spinal cord tissue was filtered through a 70 μm filter and centrifuged at 600g for 6 minutes at 4°C. The pellet was resuspended in chilled 0.1% dithiothreitol, 0.125% RNAse inhibitor, 4.2% BSA, 0.1% spermidine, and 0.1% spermine in Tris-HCl. A 50% iodixanol solution was prepared by mixing 5 parts of chilled Optiprep to 1 part 1M KCl, 1M $MgCl_2$, 1M Tricine-KOH in RNAse free water. This solution was added in equal volume to the nuclear suspension for a final 25% iodixanol. The suspension was layered above a 40% and 30% discontinuous iodixanol gradient which was then centrifuged at 10,000g for 18 minutes. The 25% iodixanol layer and approximately one third of the 30% iodixanol layer were aspirated, and the isolated sample was collected from between the 30% and 40% iodixanol layers and diluted in 2% BSA in PBS with RNAse inhibitor (RNAsin Plus, Promega N2615) for 10X RNAseq. An additional aliquot was taken and stained with DAPI for nuclei counting using a hemocytometer. During protocol optimization phase, we also performed sucrose gradient protocol with and without FACS using DAPI along with the iodixanol gradient method, for preparing 1:1 mixture of nuclei isolated from mouse brain and human spinal cord. Cross-contaminated nuclei, including doublets, were ~3% with the iodixanol, but ~27% for the sucrose gradient methods (Fig. S4).

**Library preparation and Sequencing** Nuclear isolation and snRNAseq were performed in 4 batches (TST11434, TST11456, TST11572, and TST11841). Libraries for the first two were prepared with Chromium Single Cell 3′ v3 Reagent Kits (10X Genomics, PN-1000075) according to the manufacturer's instructions. Nuclei were loaded at 1000 nuclei/μL aiming for recovery of 5000 nuclei per reaction. Libraries were sequenced on a HiSeq 2500 (Illumina) targeting 50,000 reads per nucleus. Libraries for the other two batches were prepared with Chromium Next GEM Single Cell 3′ Kit v3.1 (10X Genomics, PN-1000268) according to the manufacturer's instructions. Nuclei were loaded at 1000 nuclei/μL aiming for recovery of 5000 nuclei per reaction. Libraries were sequenced on a NovaSeq 6000 (Illumina) targeting 50,000 reads per nucleus.

**snRNAseq data processing**: CellRanger (v3.0.0) was used for demultiplexing, fastq generation from bcl, and alignment to genome (GRCh38), to generate unique molecular identifier (UMI) count tables. CellBender[11], with default parameters, was used to computationally remove potential cross-contamination among barcodes for each nucleus (hereafter “barcode”, distinguishing from Illumina index and oligo sequences for UMI). Samtools (v1.20), grep, and awk were used to parse and calculate the number of reads mapped to introns and exons for each barcode from the BAM files, which were used as a QC metric (below) (Fig. S5). Scanpy (v1.9.3) was used for handling UMI matrix and meta data for QC and clustering. Samples with high median mitochondrial percentage (%MT, >2.5%) were removed, as they also had low RIN (<3), while the reverse was not true. Nuclei with <400 or >400,000 total UMI counts, %MT of >60%, and intronic-mapped rate < 40% were removed. Barcodes originating from either VH or WSC-VH were combined per sample, and not distinguished for most of the downstream analysis.

The UMI counts were normalized by total counts per nucleus, scaled to 10,000 counts, and then log-transformed with additional pseudo-count of 1. With the normalized count table, clustering was done in a hierarchical manner. First, to identify broad cell types, cohort1 and 2 were pooled together and subjected to harmonization using harmony_integrate function (sample ID as batch, n_comps=100, n_pcs=100, theta=1, lambda=1.), then Louvain clustering was performed (n_pcs=25, n_neighbors=20, resolution=0.35) with highly-varying genes. Using known marker genes, broad cell types were identified (“level0”). To identify subtypes (“level2”), neuronal nuclei were isolated, and label transfer was performed using Seurat’s (v5.1.0) FindTransferAnchors (dims=1:30) and TransferData (dims=1:30) functions based on mouse spinal cord snRNAseq data[12]. The transferred labels were integrated into the Scanpy object, and de-novo clustering was performed as in the first step (n_pcs=50, n_neighbors=10, resolution=0.6). Louvain labels were compared to the transferred cell types, and when they match and showed consistency with the known marker genes, the transferred labels were retained. If nuclei lacked transferred label but belonged to the aforementioned cluster, the transferred label was expanded. Marker genes from previous spinal cord studies[13-15] were also cross-referenced. This process has been repeated per subclusters. When the separation between clusters were poor, initial level2 types were combined due to poor separation (e.g., DE_1-5 being DE1 and DE5 aggregated), while other level2 types were missing (e.g., VE_2). In addition, to prevent spurious identification of over-clustered cell types, cell type-to-cell type correlation of gene expressions were monitored, and when similar, they were combined into single cell type. Similar iteration was applied to glial cells. The level2 cell type identification of oligodendrocytes, OPCs, and astrocytes relied on previous publications[13,16], while microglia required deeper investigation (Fig. S9): de-novo clustering on microglia was done, and DEG of 1 vs rest using Wilcoxon rank-sum test was performed. Top DEGs were subjected to GO enrichment analysis to characterize the level2 microglia subtypes[17]. Marker genes from previous studies in ALS[18], AD[17,19,20], and cross-disease[21] were cross-referenced. Note that ventral neurons are also found in other regions as shown in Fig. S7 and 8 in Yadav et al[13], and the top genes from the ventral neurons correlate with gene expression in other regions from a previous spatial transcriptomics data[22] (Fig. S8).

**scCODA** For cell-type fractional difference between ALS and control, cohort1 and cohort2 data were subjected to scCODA (v0.1.9) separately, compared to their respective control groups. To account for over-representation of nuclei from VH, it is assumed that every spinal cord is 8mm in diameter, which gives the ratio of nuclei in two 1mm VH punches vs WSC-VH to be 3.23% ($1^2 \times 2/(8^2 - 1^2 \times 2)$). For each sample, the number of nuclei from VH were reduced to 3.23% of that from the WSC-VH, rounded to the nearest integer, and they were added together to approximate the relative number of nuclei in a WSC. All of the cell types at level0 and level2 were subjected to scCODA run, respectively, as opposed to analyzing a subset of cell types. Default parameters were used, with endothelial cells as reference, and sex and RIN as covariates. In addition to the estimated WSC counts, scCODA was also run for nuclei from VH (Supplementary Table 7).

**EWCE and MAGMA.celltyping** To identify cell types that are enriched for the expression of GWAS gene, EWCE[24] and MAGAM.celltyping[25] were used. The specificity scores were calculated using total-count normalized UMI in linear scale ("nUMI") scaled to 10,000, for control samples of cohort1 and 2 separately, at cell type level0. Genes with maximum average nUMI per cell type less than 0.5 were discarded, as specificity of those might be unreliable. EWCE (v1.6.0) was run with 10,000 permutations with default parameters. For MAGMA (v1.10), gene location files and LD reference were downloaded from the MAGMA website (https://ctg.cncr.nl/software/magma), both being GRCh37 to match that of the GWAS summary statistics. MAGMA.celltyping (v2.0.11) was used to perform linear regression of quantiles between the z-scores of genes from MAGMA and the the specificity scores. For the RNAseq data, EWCE was also applied to DEG, DSG, and gene sets from enriched pathways, with specificity score derived from control samples of cohort1. For microglia modules in hdWGCNA, genes with kME>0.7 (below) were used as gene sets for EWCE.

**Differential gene expression (DEG) analysis.** Nebula[26] was used for DEG analysis as recommended by Gagnon et al[27]. Cells were retained if they expressed at least 250 genes, while genes were included if expressed in at least three cells within the corresponding cell type. Additionally, a minimum of 3 cells per subject was required to ensure data robustness. The DEG analysis incorporated the following covariates: age, sex, RIN, sequencing batches, mitochondrial gene percentage, intronic mapped rate, and total UMI counts. TDP43 pathology score regression is also performed using Nebula, with the same set of covariates as in disease vs control. Note that the pathology score is at subject level, not at single-cell level. Pathway analysis was done as in RNAseq Data Analysis.

**Countsplit:** To avoid "double-dipping" problem in differential gene expression analysis between subtypes of cells and hdWGCNA module preservation analysis (below), we applied Countsplit[28] with a Poisson model (overdisps=NULL, folds=2), using epsilon=c(0.3,0.7) where "0.3" for 30% down-sampling of UMI counts for the training set, and "0.7" for the test set. The training set was used for (1) reclustering of microglia subtypes, and (2) building the consensus module in hdWGCNA (below). The test set was used for (1) DEG analysis of between-subtype of microglia (e.g., MG1 vs MG0), and (2) module preservation calculation between ALS types (below). Note that for microglia, MG2 was not detected as a separate entity in the reclustering, but mostly became part of MG1.

**Translatability analysis across ALS subtypes – hdWGCNA** To assess the preserved gene-gene network structures across ALS subtypes, we constructed co-expression networks for each ALS subtype using the single-nucleus weighted gene co-expression network analysis (hdWGCNA) approach[29], stratified by cell type. First, the network was constructed using the "training set" for each cell type and ALS type. We then constructed a consensus co-expression network by computing the TOM across the three ALS subtypes: sALS, C9, and SOD1. Finally, we evaluated preservation of the modules (Zsummary statistics[30] ) between ALS types, using the "test set". The detailed procedure is outlined below.

**Construction of Co-Expression Networks** Single-cell gene expression datasets are inherently sparse, with a predominance of zero-valued entries. This sparsity disrupts correlation coefficient calculations in co-expression analyses. To mitigate this issue, we constructed meta-cells by aggregating single-cell transcriptomes through a bootstrapped approach. Specifically, for each meta-cell, a randomly selected nucleus was aggregated with its 50 nearest neighboring nuclei. To ensure the uniqueness of each meta-cell, we restricted the overlap between meta-cells to a maximum of 50%. For less abundant cell types, such as motor neurons, we relaxed this criterion, allowing up to 80% overlap. We then applied the standard WGCNA pipeline to compute TOM for each ALS subtype, stratified by cell type.

**Consensus Module Identification and Zsummary calculation** Using the training set, we computed TOM for each ALS subtype, stratified by cell type. The consensus TOM matrix for each cell type was obtained by calculating the element-wise median across the three ALS subtype-specific TOM matrices (sALS, C9, and SOD1). Consensus modules were identified as dendrogram branches in the consensus TOM using the Dynamic Tree Cut algorithm. We employed weighted correlation with individual sample weights, as described above, and constructed a "signed hybrid" network, wherein only positively correlated genes were considered connected. The module merging threshold was optimized to maximize the correlation between module eigengenes and disease status. To evaluate module preservation across ALS subtypes, we calculated the Zsummary statistic using the modulePreservation function in WGCNA. In WGCNA, module eigengene (ME) is the first principal component of the module, which can be considered as a meta-gene that represents the module. The correlation between ME and disease status – ALS=1, control=0 – was calculated, and further analysis was performed for the modules with high Zsummary statistic and consistent and strong ME-to-disease correlation. Pathway analysis was done similar to RNAseq Data Analysis, except for including genes with kME > 0.7 for each module, and used directional information from differential expression analysis, even for those with Q>0.05.


1 Dobin, A. & Gingeras, T. R. Optimizing RNA-Seq Mapping with STAR. *Methods Mol Biol* **1415**, 245-262 (2016). https://doi.org:10.1007/978-1-4939-3572-7_13
2 Li, B. & Dewey, C. N. RSEM: accurate transcript quantification from RNA-Seq data with or without a reference genome. *BMC Bioinformatics* **12**, 323 (2011). https://doi.org:10.1186/1471-2105-12-323
3 DeLuca, D. S. *et al.* RNA-SeQC: RNA-seq metrics for quality control and process optimization. *Bioinformatics* **28**, 1530-1532 (2012). https://doi.org:10.1093/bioinformatics/bts196
4 Love, M. I., Huber, W. & Anders, S. Moderated estimation of fold change and dispersion for RNA-seq data with DESeq2. *Genome Biol* **15**, 550 (2014). https://doi.org:10.1186/s13059-014-0550-8
5 Zhu, J. *et al.* RNASequest: An End-to-End Reproducible RNAseq Data Analysis and Publishing Framework. *J Mol Biol* **435**, 168017 (2023). https://doi.org:10.1016/j.jmb.2023.168017
6 Zhu, A., Ibrahim, J. G. & Love, M. I. Heavy-tailed prior distributions for sequence count data: removing the noise and preserving large differences. *Bioinformatics* **35**, 2084-2092 (2019). https://doi.org:10.1093/bioinformatics/bty895
7 Shen, S. *et al.* rMATS: robust and flexible detection of differential alternative splicing from replicate RNA-Seq data. *Proc Natl Acad Sci U S A* **111**, E5593-5601 (2014). https://doi.org:10.1073/pnas.1419161111
8 Li, Y. I. *et al.* Annotation-free quantification of RNA splicing using LeafCutter. *Nat Genet* **50**, 151-158 (2018). https://doi.org:10.1038/s41588-017-0004-9
9 Vaquero-Garcia, J. *et al.* RNA splicing analysis using heterogeneous and large RNA-seq datasets. *Nat Commun* **14**, 1230 (2023). https://doi.org:10.1038/s41467-023-36585-y

10 Hrvatin, S. *et al.* A scalable platform for the development of cell-type-specific viral drivers. *Elife* **8** (2019). https://doi.org:10.7554/eLife.48089
11 Fleming, S. J. *et al.* Unsupervised removal of systematic background noise from droplet-based single-cell experiments using CellBender. *Nat Methods* **20**, 1323-1335 (2023). https://doi.org:10.1038/s41592-023-01943-7
12 Sathyamurthy, A. *et al.* Massively Parallel Single Nucleus Transcriptional Profiling Defines Spinal Cord Neurons and Their Activity during Behavior. *Cell Rep* **22**, 2216-2225 (2018). https://doi.org:10.1016/j.celrep.2018.02.003
13 Yadav, A. *et al.* A cellular taxonomy of the adult human spinal cord. *Neuron* **111**, 328-344 e327 (2023). https://doi.org:10.1016/j.neuron.2023.01.007
14 Blum, J. A. *et al.* Single-cell transcriptomic analysis of the adult mouse spinal cord reveals molecular diversity of autonomic and skeletal motor neurons. *Nat Neurosci* **24**, 572-583 (2021). https://doi.org:10.1038/s41593-020-00795-0
15 Abdel-Khalik, J. *et al.* Defective cholesterol metabolism in amyotrophic lateral sclerosis. *J Lipid Res* **58**, 267-278 (2017). https://doi.org:10.1194/jlr.P071639
16 Seeker, L. A. *et al.* Brain matters: unveiling the distinct contributions of region, age, and sex to glia diversity and CNS function. *Acta Neuropathol Commun* **11**, 84 (2023). https://doi.org:10.1186/s40478-023-01568-z
17 Sun, N. *et al.* Human microglial state dynamics in Alzheimer's disease progression. *Cell* **186**, 4386-4403 e4329 (2023). https://doi.org:10.1016/j.cell.2023.08.037
18 Limone, F. *et al.* Single-nucleus sequencing reveals enriched expression of genetic risk factors in extratelencephalic neurons sensitive to degeneration in ALS. *Nat Aging* **4**, 984-997 (2024). https://doi.org:10.1038/s43587-024-00640-0
19 Prater, K. E. *et al.* Human microglia show unique transcriptional changes in Alzheimer's disease. *Nat Aging* **3**, 894-907 (2023). https://doi.org:10.1038/s43587-023-00424-y
20 Keren-Shaul, H. *et al.* A Unique Microglia Type Associated with Restricting Development of Alzheimer's Disease. *Cell* **169**, 1276-1290 e1217 (2017). https://doi.org:10.1016/j.cell.2017.05.018
21 Tuddenham, J. F. *et al.* A cross-disease resource of living human microglia identifies disease-enriched subsets and tool compounds recapitulating microglial states. *Nat Neurosci* **27**, 2521-2537 (2024). https://doi.org:10.1038/s41593-024-01764-7
22 Maniatis, S. *et al.* Spatiotemporal dynamics of molecular pathology in amyotrophic lateral sclerosis. *Science* **364**, 89-93 (2019). https://doi.org:10.1126/science.aav9776
23 Buttner, M., Ostner, J., Muller, C. L., Theis, F. J. & Schubert, B. scCODA is a Bayesian model for compositional single-cell data analysis. *Nat Commun* **12**, 6876 (2021). https://doi.org:10.1038/s41467-021-27150-6
24 Skene, N. G. & Grant, S. G. Identification of Vulnerable Cell Types in Major Brain Disorders Using Single Cell Transcriptomes and Expression Weighted Cell Type Enrichment. *Front Neurosci* **10**, 16 (2016). https://doi.org:10.3389/fnins.2016.00016
25 Skene, N. G. *et al.* Genetic identification of brain cell types underlying schizophrenia. *Nat Genet* **50**, 825-833 (2018). https://doi.org:10.1038/s41588-018-0129-5
26 He, L. *et al.* NEBULA is a fast negative binomial mixed model for differential or co-expression analysis of large-scale multi-subject single-cell data. *Commun Biol* **4**, 629 (2021). https://doi.org:10.1038/s42003-021-02146-6
27 Gagnon, J. *et al.* Recommendations of scRNA-seq Differential Gene Expression Analysis Based on Comprehensive Benchmarking. *Life (Basel)* **12** (2022). https://doi.org:10.3390/life12060850

28 Neufeld, A., Gao, L. L., Popp, J., Battle, A. & Witten, D. Inference after latent variable estimation for single-cell RNA sequencing data. *Biostatistics* **25**, 270-287 (2023). https://doi.org:10.1093/biostatistics/kxac047

29 Morabito, S., Reese, F., Rahimzadeh, N., Miyoshi, E. & Swarup, V. hdWGCNA identifies co-expression networks in high-dimensional transcriptomics data. *Cell Rep Methods* **3**, 100498 (2023). https://doi.org:10.1016/j.crmeth.2023.100498

30 Langfelder, P., Luo, R., Oldham, M. C. & Horvath, S. Is my network module preserved and reproducible? *PLoS Comput Biol* **7**, e1001057 (2011). https://doi.org:10.1371/journal.pcbi.1001057
